# Rapid Single-Shot Synthesis of the 217 Amino Acid-Long N-Terminal Domain of Pyocin S2

**DOI:** 10.1101/2022.11.17.516969

**Authors:** Azin Saebi, Joseph S. Brown, Victoria M. Marando, Nina Hartrampf, Nicole M. Chumbler, Stephanie Hanna, Mackenzie Poskus, Andrei Loas, Laura L. Kiessling, Deborah T. Hung, Bradley L. Pentelute

**Author notes:** These authors contributed equally (Azin Saebi, Joseph S. Brown). Department of Chemistry, University of Zurich, 8057 Zurich, Switzerland (Nina Hartrampf).

## Abstract

The impermeable outer membrane of *Pseudomonas aeruginosa* is bypassed by antibacterial proteins known as S-type pyocins. Because of their properties, pyocins are investigated as a potential new class of antimicrobials against *Pseudomonas* infections. Their production and modification, however, remains challenging. To address this limitation, we employed automated fast-flow peptide synthesis (AFPS) for the rapid production of a pyocin S2 import domain. The N-terminal domain sequence (PyS2^NTD^) was synthesized in under 10 hours and purified to yield milligrams quantities of the desired product. To our knowledge, the 217 amino acid sequence of PyS2^NTD^ is among the longest peptides produced from a “single-shot” synthesis, i.e., made in a single stepwise route without the use of ligation techniques. Biophysical characterization of the PyS2^NTD^ with circular dichroism was consistent with the literature reports. Fluorescently labeled PyS2^NTD^ binds to *P. aeruginosa* expressing the cognate ferripyoverdine receptor (FpvA) and is taken up into the periplasm. This selective uptake was validated with confocal and super resolution microscopy, flow cytometry, and fluorescence recovery after photobleaching (FRAP). These modified, synthetic S-type pyocins domains can be used to probe import mechanisms of *P. aeruginosa* and leveraged to develop selective antimicrobial agents that bypass the outer membrane.

## Introduction

Chemical synthesis is a powerful approach to access proteins and study their function. This strategy sidesteps biological expression bottlenecks to yield proteins that would be otherwise difficult to obtain (e.g., unstable, or toxic proteins).^1-3^ Chemical protein synthesis also allows for unmatched flexibility in the incorporation of noncanonical residues. Thus, chemical protein synthesis can provide access to proteins with a plethora of chemical modifications to optimize their properties and activity, including efficacy, bioavailability, and half-life.^4,5^ Most methods towards chemical protein synthesis rely on solid-phase peptide synthesis (SPPS).^6^ This process involves synthesis of the peptide backbone, cleavage, and deprotection of the side chains followed by purification and folding to yield a functional protein. While this method is generally successful, the accumulation of side products during SPPS can limit the purity and isolated yield of the desired full-length peptide. To maintain a straightforward purification, peptide synthesis is often limited to a chain length of 30-50 residues, which is significantly smaller than the average length of a single-domain protein.^7-10^ Chemical protein synthesis has therefore relied heavily on ligation of shorter peptide fragments prepared in convergent SPPS. Most commonly, native chemical ligation (NCL) or α-ketoacid-hydroxylamine ligation (KAHA) strategies are employed to this end.^11,12^

Ligation strategies have delivered an impressive number of proteins, with the primary challenges being the production of multiple fragments and efficacy of the ligation assembly steps. These approaches have yielded proteins including mirror-image Pfu DNA polymerase with 775 amino acids,^13^ tetra-ubiquitin-α-globin with 472 amino acids,^14^ the Lys11/Lys48-branched hexaubiquitin chain with 456 amino acids,^15^ P2 DNA polymerase IV with 358 amino acids,^16^ DapA with 312 amino acids,^17^ and the F-ATPase subunit γ with 286 residues.^18^

These studies are a testament to the power of chemical protein synthesis; however, ligation strategies remain limited by the time to produce and assemble multiple fragments. Fragments can have poor solubility properties compared to the full intact protein, which has given rise to the design and use of solubility tags in ligation.^19,20^ As protein length increases toward single-domain proteins and beyond, the challenge of chemical synthesis and ligation can become demanding and require multiple approaches to be explored to identify high yielding routes. Approaches including possible mutations to accommodate ligation sites, sequential deprotection of cysteines, and/or desulfurization after ligation can be considered. Moreover, purification of individual fragments or ligated fragments can be time-consuming and potentially introduce yield limiting steps even before approaching final protein folding and biopurification.^21^

The rapid and routine access to large synthetic proteins can be streamlined by technological and chemical advances to generate longer peptide sequences, thus decreasing the number of ligations required. Significant time-savings could be achieved if protein synthesis were completed in a “single-shot,” wherein the full-length polypeptide chain was produced in a single SPPS effort at sufficient purity and yield. This approach would minimize intermediate handling steps, reactions, and purifications, prior to folding. However, single-shot peptide synthesis of long sequences has historically been limited by steric hindrance of protecting groups, formation of secondary structures, and intra- and/or intermolecular aggregation on resin. All of these barriers interfere with synthesis, reducing purity and yield.^9,22-24^ Approaches to improve SPPS by focusing on overcoming on-resin aggregation, including use of backbone protected monomers,^25,26^ alternative solvents,^27^ and microwave heating.^28,29^ If known, structural information can also inform the optimization of the protein sequence to improve synthesis efficiency.^30-32^ Overall, these strategies have led to the robust synthesis of difficult peptides with decreased synthesis time and increased crude peptide purity.

Flow chemistry has advanced SPPS by improving the synthesis rate and quality of long peptides through rich history of development.^33-38^ Our group recently implemented automated fast-flow peptide synthesis (AFPS), a technology for rapid synthesis of peptides in flow.^39^ Rigorous optimization of reaction parameters including use of amine-free *N,N*-dimethylformamide (DMF), increased amino acid concentration to 0.4 M, and increased reagent activation and coupling temperature to 90 °C led to improved results. The use of formic acid as an additive for fluorenylmethyloxycarbonyl (Fmoc) deprotection,^40^ and optimization of cysteine and histidine couplings led to reduction in aspartimide formation and epimerization, respectively. These changes thereby minimize some of the known challenges of peptide synthesis at elevated temperature. Further residue-specific optimization for coupling agents resulted in the establishment of a general protocol for AFPS.^41,42^ This protocol led to the successful syntheses of single-domain proteins in a single-shot approach, including murine double minute 2 protein domain (MDM2 domain, 127 AA), lysozyme (129 AA), fibroblast growth factor 1 (FGF1, 140 AA), and sortase A* (164 AA).

These initial successes prompted us to apply this optimized AFPS protocol to produce a protein over 200 residues long in a single-shot synthesis. Our objective was to assess whether this approach could yield workable amounts (>1 mg) of biologically active material for functional studies. To this end, we describe here the single-shot synthesis of the 217-mer N-terminal receptor binding domain of the bacteriocin pyocin S2 (PyS2^NTD^) produced by the Gram-negative bacterium *Pseudomonas aeruginosa*.^43^ Full-length pyocin S2 mimics the pyoverdine siderophore, binds to the ferripyoverdine Type 1 receptor (FpvA), and crosses both the outer and inner membranes to act as a toxic nuclease in the cytoplasm of *P. aeruginosa*.^44^ While the C-terminal region of the protein displays nuclease activity, the N-terminal domain (PyS2^NTD^) is involved in facilitating pyocin S2 binding and internalization to the periplasm, confirmed in literature^45^ and by our study here with synthetically produced PyS2^NTD^. Given that the low permeability of the outer membrane of *P. aeruginosa* is a major contributor to its resistance against antibiotics,^46,47^ we envision the production of PyS2^NTD^ will enable the development of novel antipseudomonal therapeutics and further improve the understanding of pyocin transport.^48,49^

## Results and Discussion

### AFPS enables rapid single-shot synthesis of PyS2^NTD^

The N-terminal domain of pyocin S2 (PyS2^NTD^, Q06584, M1-K209) has been previously produced using recombinant DNA technology and shown to translocate across the outer membrane of *P. aeruginosa* via its native FpvA receptor.^44,45^ We hypothesized this protein fragment could be feasibly produced in a single-shot synthesis, rather than through ligation (Figure 1A) due to advancements made in automated flow technology for peptide and protein synthesis. Structural and functional data in the literature provide a direct comparison to understand and verify the integrity and function of synthetic PyS2^NTD^.^45^ To minimize any disruption in the ability of the N-terminal domain to undergo internalization, a cysteine handle for site-selective bioconjugation was incorporated near the C-terminus (Cys216) of the PyS2^NTD^ sequence. Furthermore, to reduce the possibility of methionine oxidation side products, methionine residues were mutated to norleucine (Nle),^30^ resulting in a synthetic PyS2^NTD^ construct 217 residues in length (Figure 1B).

**Figure 1.**
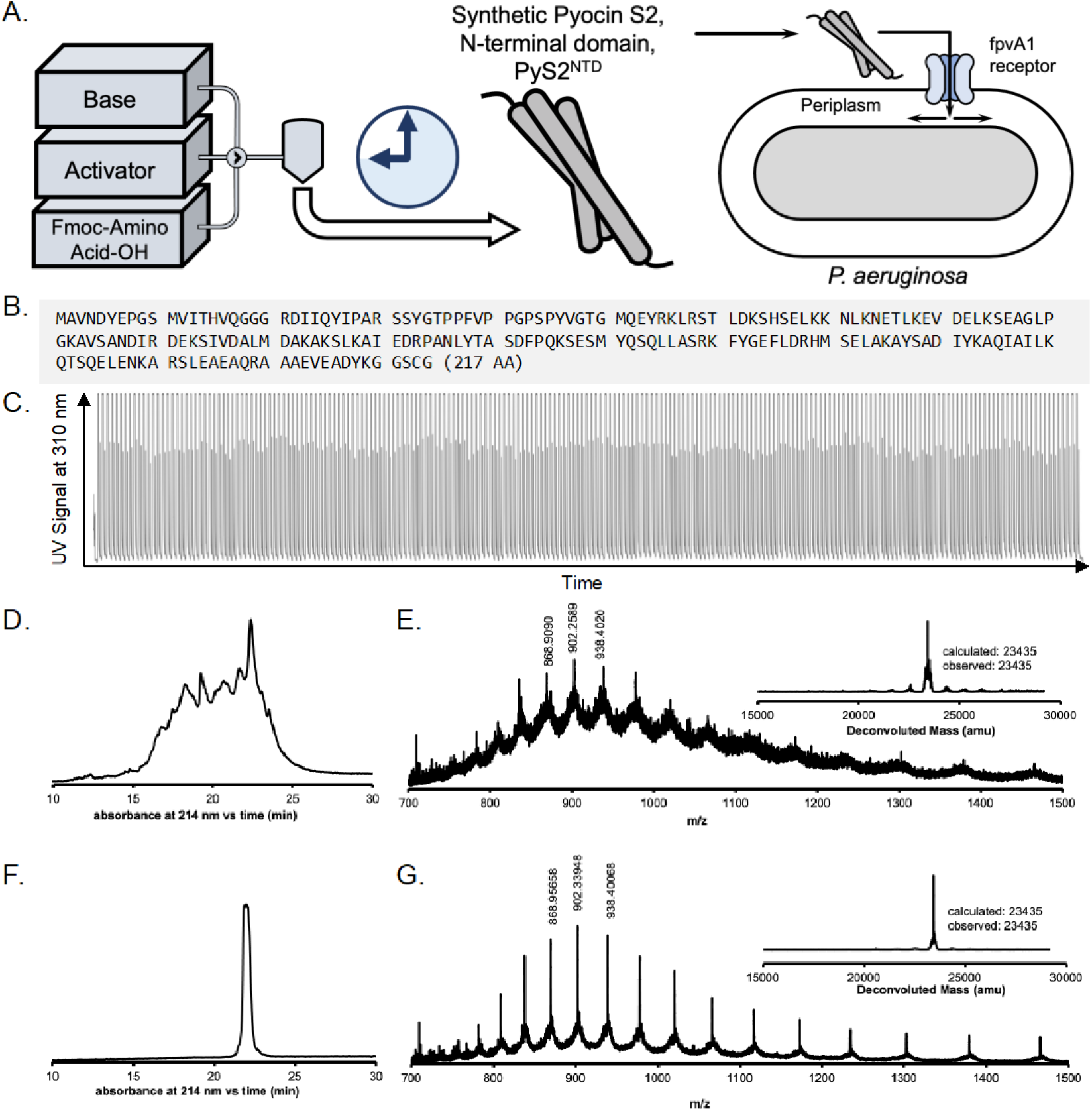
PyS2^NTD^ was rapidly synthesized in 9.2 hours using automated fast-flow peptide synthesis (AFPS). **A**. Schematic representation of the automated flow synthesis and biological function of PyS2^NTD^. **B**. The sequence of PyS2^NTD^ with cysteine (Cys216) added for bioconjugation. **C**. The UV absorbance trace at 310 nm from flow synthesis showed the Fmoc-deprotection peaks remained comparable in height and width throughout the synthesis. Coupling steps are also shown and produce a saturated signal, just before each Fmoc-deprotection peak. **D**. Analytical reverse-phase high-performance liquid chromatography (RP-HPLC) from absorbance at 214 nm and **E**. liquid chromatography–mass spectrometry (LC-MS) mass spectrum of crude PyS2^NTD^ with deconvoluted mass spectrum inset. **F**. RP-HPLC of purified PyS2^NTD^ with absorbance at 214 nm and **G**. LC-MS mass spectrum of the purified PyS2^NTD^ with deconvoluted mass spectrum inset. For additional information on synthesis, cleavage, and purification conditions see Supporting Information SI 2.0. through SI SI 7.0.

PyS2^NTD^ was synthesized in an automated flow sequence of *in situ* amino acid activation, residue incorporation via amide coupling on solid-phase H-Rink ChemMatrix resin, and deprotection of the newly incorporated residue (Supporting Information SI 4.0.). The efficiency of residue incorporation was monitored in real time via an in-line UV-Vis absorbance detector. At 310 nm, the release of fluorene-containing compounds was observed, allowing for quantification of the deprotection step as a measure of residue incorporation.^41,42,50^ Any irregularities, such as a decrease in deprotection peak height, could be indicative of aggregation of the nascent chain.^23,50^ Analysis of the peak area resulting from the amine deprotection during the synthesis of PyS2^NTD^ suggested no major aggregation events (Figure 1C, Figure S1). This synthesis protocol was previously developed for the production of shorter sequences and used here with no further optimization or modification, supporting its robustness and potential general applicability.^42^ Synthesis of PyS2^NTD^ proceeded for 9.2 hours with 434 steps, where each residue was incorporated in approximately 2.5 minutes.

After synthesis, the crude PyS2^NTD^ clearly exhibited the expected mass corresponding to the intact protein, and preparative reverse-phase purification was able to isolate >10 mg of the unfolded PyS2^NTD^ polypeptide. Cleavage provided 197 mg of crude PyS2^NTD^ as the trifluoroacetate salt from 80 mg of resin after lyophilization. The synthesis quality of the crude PyS2^NTD^ was examined by analytical liquid chromatography and liquid chromatography-mass spectrometry (LC-MS, Figures S2-S6, Supporting Information SI 5.0.). Analytical reverse-phase chromatography of the crude PyS2^NTD^ provided a quality assessment of potential synthesis issues including truncations, deletions, and isomers. Though the PyS2^NTD^ exhibited multiple peaks (Figure 1D), the presence of some distinct peaks indicated that optimized preparative chromatography could potentially yield pure fractions. Using LC-MS, the crude synthesis quality was evaluated to determine if the calculated mass of the fully assembled N-terminal domain was present. When integrating the entire mass chromatogram peak from the crude PyS2^NTD^, a charge-state series of ions were observed in the raw mass chromatogram (Figure 1E). From this raw chromatogram, a deconvoluted mass spectrum clearly exhibited the expected calculated molecular weight of the fully assembled PyS2^NTD^ (Figure 1E inset). PyS2^NTD^ was purified by preparative reverse-phase high-performance liquid chromatography (RP-HPLC, Figures S8-S17, Supporting Information SI 6.0. and 7.0.). A preparative-grade C18 column was used at elevated temperature (60 °C) to isolate 10.3 mg of pure PyS2^NTD^ as the trifluoroacetate salt. LC-MS was again used to confirm the molecular weight of the product and analytical HPLC confirmed isolation of PyS2^NTD^ from multiple side products (Figures S18-S22).

### Folded synthetic PyS2^NTD^ possesses the structure and melting temperature

After reverse-phase preparative HPLC purification, PyS2^NTD^ was folded from the polypeptide chain into its native, globular folded structure in high yield (84%). Folding strategies typically include dilution, dialysis, and solid phase-assisted refolding.^21,51^ Generally, solubilization and denaturation of the protein in a high concentration of a chaotrope is followed by controlled removal of the chaotrope to trigger folding. We utilized a dilution-based method for PyS2^NTD^ involving size exclusion chromatography (SEC). SEC-based folding was achieved by simultaneously desalting the PyS2^NTD^ from 6 M guanidine hydrochloride upon sample injection as well as purifying the protein from non-folded aggregates on the column. As PyS2^NTD^ does not include any intramolecular disulfide bonds, no oxidative/reductive buffer potential was required (Figure 2B and Supporting Information SI 8.0.). SEC-folded PyS2^NTD^ yielded a product with excellent purity (Figure 2C). The charge state series observed in the LC-MS spectrum was better resolved, indicating improved purity of the synthetic PyS2^NTD^ protein (Figure 2D). With this method, pure PyS2^NTD^ fractions did not need to be concentrated, yielding 0.73 mg (31 nmol) PyS2^NTD^ from 1.0 mg injected (37 nmol).

**Figure 2.**
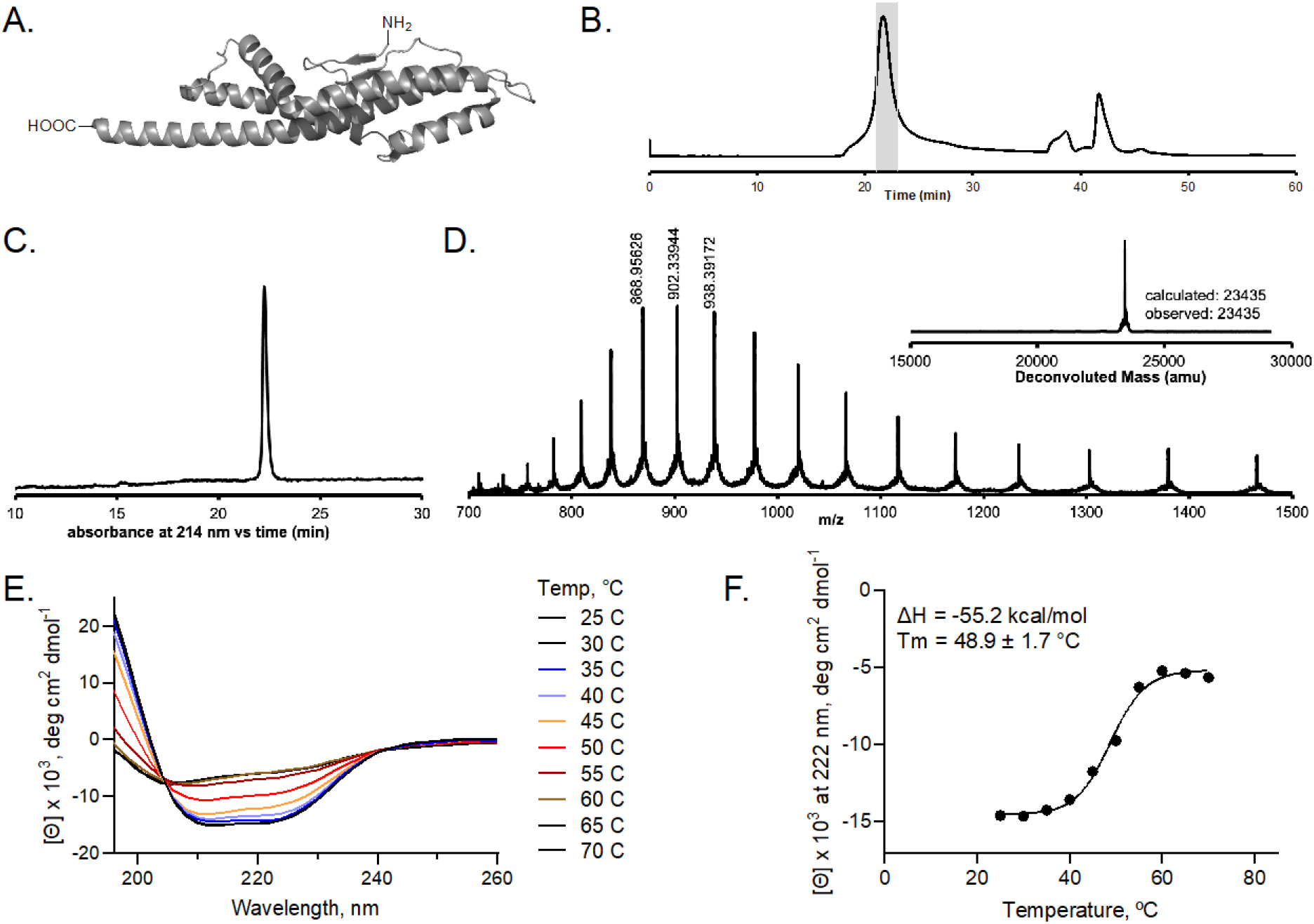
Biophysical characterization of synthetic PyS2^NTD^ after SEC-based folding. **A**. Crystal structure of PyS2^NTD^ (residues 11-205, PDB: 5ODW) displays primarily α-helices and a short antiparallel β-hairpin. **B**. Chromatogram of size exclusion chromatography (SEC) purification of PyS2^NTD^, showing absorbance at 214 nm, where the fractions containing the protein (shaded in gray) were pooled to yield 0.73 mg of material (84% yield). **C**. RP-HPLC of folded PyS2^NTD^ showing absorbance at 214 nm and **D**. LC-MS of the folded PyS2^NTD^ with deconvoluted mass spectrum inset. **E**. Far-UV circular dichroism (CD) spectrum of the purified synthetic PyS2^NTD^. **F**. Thermal melt curve for synthetic PyS2^NTD^ in the temperature range of 25–70 °C. The melting temperature (T_m_) was determined to be 49 ± 2 °C (Mean ± 95% CI), similar to the 50 °C value reported in literature.^45^ See Supporting Information SI 8.0. through 10.0. for details on folding, purification, and CD.

The secondary structure of the synthetic PyS2^NTD^ was examined using circular dichroism (CD). The resulting spectra were consistent with the reported recombinant material.^52^ The structure of recombinant PyS2^NTD^ complexed with its receptor FpvA determined by x-ray crystallography (PDB: 5ODW, PyS2 shown in Figure 3A)^45^ indicates that the domain is composed of five α-helices and a short antiparallel β-hairpin, with the α-helices comprising >70% of the sequence. The CD of PyS2^NTD^ has a mean residue ellipticity (MRE) signal with a local minimum at 222 nm close in depth to an absolute minimum at 208 nm (Supporting Information SI 10.0.). The MRE increased with absolute maximum at the shortest wavelength measured at 195 nm. MRE at 208 nm (negative), 222 nm (negative), and 193 nm (positive) are characteristic signatures of α-helical proteins. These signatures suggested that other structures are present with absorbance at lower wavelengths, consistent with the loop regions and the antiparallel β-sheet structures in PyS2^NTD^.

**Figure 3.**
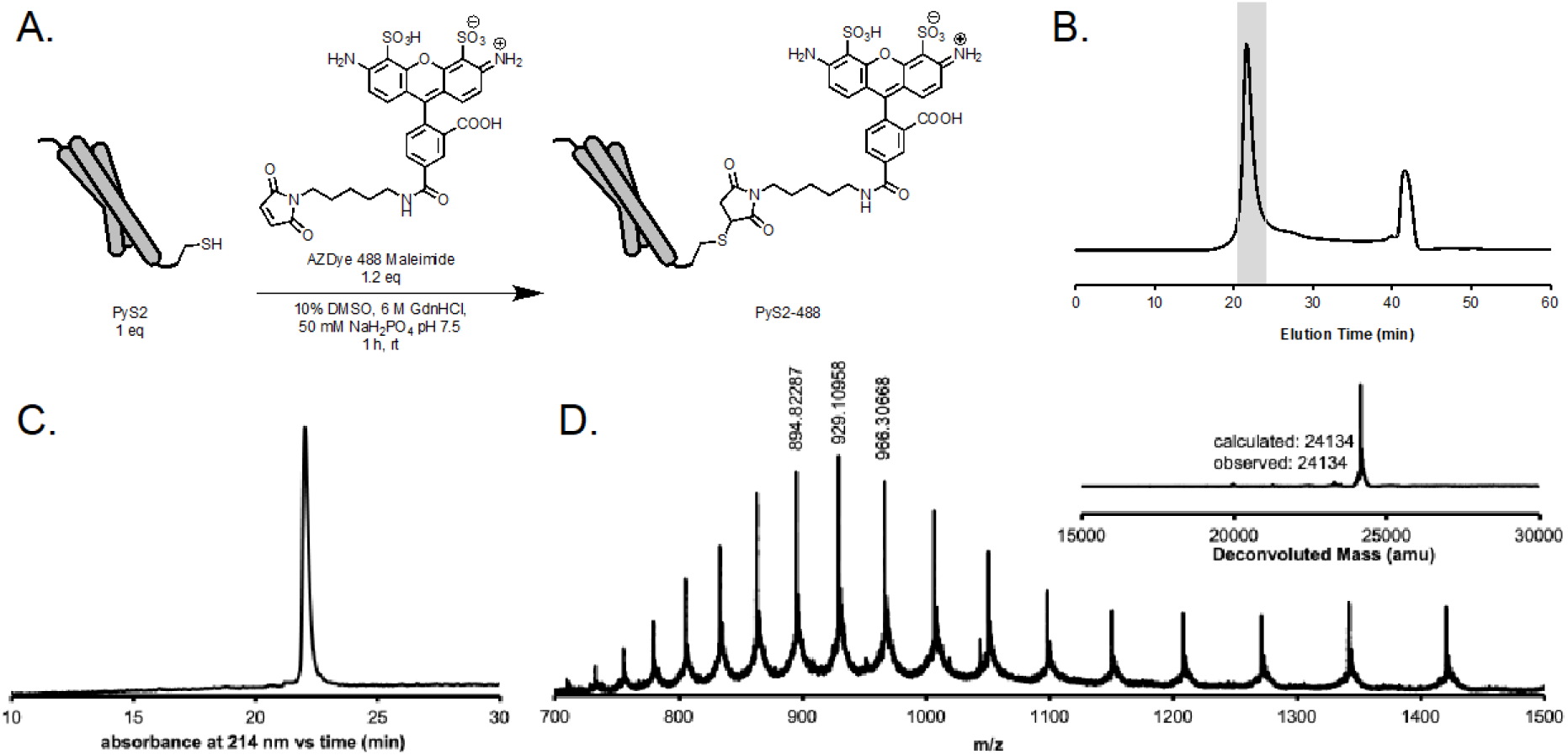
Synthetic PyS2^NTD^ was conjugated to fluorophore AZDye 488 and folded. **A**. Conjugation and folding scheme of PyS2^NTD^. **B**. Chromatogram of size exclusion chromatography (SEC) purification of PyS2-488, showing fluorescence signal with excitation at 280 nm and emission at 325 nm, where the fractions containing the protein (shaded in gray) were pooled to yield 234 °g of material corresponding to 24% combined reaction and folding recovery yield. **C**. RP-HPLC of folded PyS2-488 conjugate showing absorbance at 214 nm and **D**. LC-MS of the folded PyS2-488 conjugate with deconvoluted mass spectrum inset. See Supporting Information SI 11.0. and 12.0. for more on conjugation and purification.

To further assess the structural integrity of the synthetic PyS2^NTD^, we determined its melting temperature (T_m_), which matched that of the reported value of the recombinant material (Figure 3E-F and Supporting Information SI 10.0.).^53^ The MRE response weakened with increased temperature, and the inflection of the transition marked the measured PyS2^NTD^ melting temperature (T_m_, Figure 2F). Melting was complete with no appreciable change in CD signal observed beyond 60 °C. The unfolded protein displayed low ellipticity with an absolute minimum near 205 nm. Overall, structural analysis of PyS2^NTD^ demonstrates that the sequence forms a predominantly α-helical structure with a melting temperature calculated to be 49 ± 2 °C (Mean ± 95% CI), consistent with the literature value (50 °C) for the recombinant protein.^45^

### Site specific PyS2^NTD^ labeling with AZDye 488 via cysteine-maleimide conjugation

To functionally characterize the synthetic PyS2^NTD^, we prepared a synthetic PyS2^NTD^-fluorophore conjugate using the installed cysteine residue. Fluorescence microscopy has been previously used to assess the import of recombinant PyS2^NTD^ into live *P. aeruginosa* cells.^45^ Although folded synthetic PyS2^NTD^ can be labeled under aqueous conditions, we sought to combine the conjugation and folding processes to streamline the procedure and maximize PyS2^NTD^ yield. Accordingly, HPLC-purified synthetic PyS2^NTD^ was simultaneously denatured and labeled with AZDye 488 (1.2 equiv) via cysteine-maleimide conjugation (Figure 4A). The reaction was monitored by LC-MS (Figure 4B) for the expected mass shift of +698 Da, corresponding to the addition of the AZDye to the PyS2^NTD^. The labeling and folding steps afforded the desired folded PyS2-488, corresponding to 24% isolated yield (234 ug from 1.17 mg reacted, Figure 4C). The purity of the PyS2-488 post-folding and conjugation was high as evaluated by analytical RP-HPLC and LC-MS (Figure 4C, D, E, see Supporting Information SI 11.0. and 12.0.).

**Figure 4.**
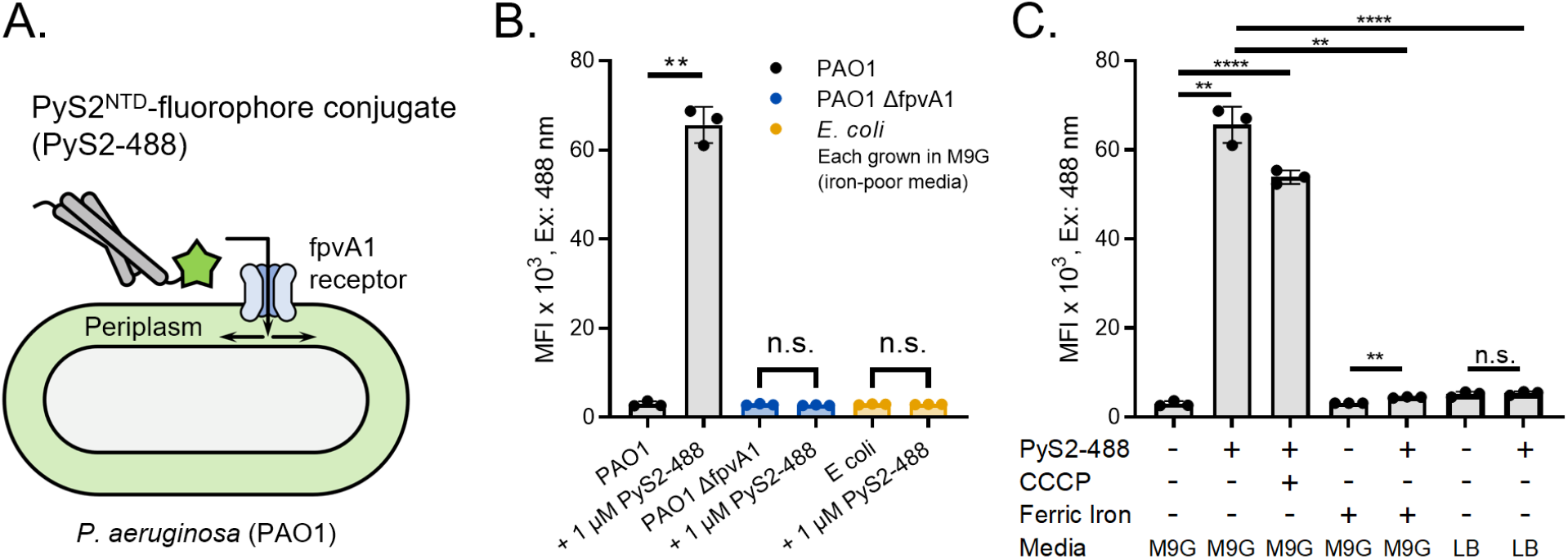
Synthetic PyS2^NTD^-488 binds *P. aeruginosa* with expected species- and receptor-specificity. **A**. The FpvA-dependent internalization of PyS2^NTD^-fluorophore conjugate can be monitored by the emitted fluorescence signal via flow cytometry. **B**. Flow cytometry indicates species specificity staining of PyS2-488 with only PAO1 positively stained in comparison to control (*p* = 0.0011), whereas knockout strain PAO1 Δ*fpvA* and *E. coli* (ATCC 25922) showed no staining when grown in iron-poor minimal M9 Glucose (M9G) media (*p* = 0.297 and *p* = 0.837, respectively). **C**. Growth and staining conditions known to affect the abundance and function of FpvA further confirmed that PyS2-488 is binding to FpvA. Positive staining is observed in minimal M9G compared to control (*p* = 0.0011). The addition of the CCCP internalization inhibitor (100 µM pretreatment) minimally affected PyS2-488 staining as it only affects periplasmic internalization and accumulation, leaving FpvA abundance and thus PyS2-488 staining unaffected (*p* < 0.0001 compared to control PAO1). Limited staining was observed if ferric iron is added to M9G or iron-rich media (Luria Broth, LB) is used because the iron scavenging function of the FpvA is less critical (*p* = 0.0014 and < 0.0001 compared to PAO1 in M9G, respectively).^57^ Gating strategy, histograms, and representative raw cytometry plots can be seen in the Supporting Information SI 13.3. All experimental conditions were completed with *n* = 3 biological replicates with individual data points shown and all statistical t-tests shown are paired t-test, two-tailed with Welch’s correction with *p* ≤ 0.01, 0.001, 0.0001, represented as **, ***, and **** respectively.

### PyS2^NTD^-fluorophore conjugate binds and internalizes into the *P. aeruginosa* periplasm

The uptake PyS2^NTD^-AZDye488 conjugate (PyS2-488) into *P. aeruginosa* was then evaluated using flow cytometry and fluorescence microscopy (Figure 4A). Natively, PyS2 binds to the ferripyoverdine receptor (FpvA) with high affinity (dissociation constant, *K*_D_ < 1 nM)^45^ and undergoes internalization through a process in which the proton motive force coupled to the inner membrane protein TonB1 facilitates transport after FpvA binding. Thus, PyS2-488 should demonstrate species specificity for *P. aeruginosa* and receptor specificity for FpvA1. PAO1 was used as an *fpvA*-positive strain, with *fpvA*-negative controls including *E. coli* (ATCC 25922) and a PAO1 FpvA null (Δ*fpvA* deletion, see Supporting Information SI 13.1.). Treatment of the bacteria with PyS2-488 revealed species and receptor-specific staining with PAO1 only demonstrating a positive signal (Figure 4B, Supporting Information SI 13.0.).

Additional media conditions were examined to further test the receptor binding specificity and internalization of PyS2-488. The regulation of FpvA receptor expression depends on the bacterial environment: iron-poor conditions promote the production of the ferripyoverdine iron scavenger and its cognate receptor FpvA.^54^ Thus, the PAO1 and PAO1 Δ*fpvA* were grown in either a minimal media M9 Glucose (M9G) containing no iron or M9G supplemented with 17 µM ferric (+3) iron chloride to replicate iron-rich conditions (e.g., Luria broth).^55,56^ For control experiments, the proton motive force was diminished by the pre-treatment of PAO1 with protonophore carbonyl cyanide m-chlorophenyl hydrazone (CCCP). The use of CCCP thus inhibits PyS2-488 internalization, while leaving FpvA binding unaffected. Treatment of the bacteria with PyS2-488 revealed staining consistent with the environment- and proton motive force-dependent presence of the FpvA-TonB transport system. Specifically, the presence of ferric iron in the growth media ablated PyS2-488 staining, likely because of the lack of FpvA expression and survival dependence on iron-scavenging. CCCP had a minimal effect to reduce the strong staining of PyS2-488, likely because FpvA binding was uninhibited, but internalization and accumulation of PyS2-488 in the periplasm was slowed.

To evaluate FpvA-based binding and internalization of PyS2-488 further, fluorescence microscopy techniques were used to assess periplasmic localization of PyS2-488 of the positively stained samples (PAO1 and PAO1 + the internalization inhibitor CCCP). PAO1 cells stained with PyS2-488 are expected to achieve periplasmic internalization.^45^ However, the PyS2-488-stained PAO1 cells pretreated with the internalization inhibitor CCCP should only show outer membrane localization because CCCP diminishes the proton motor force utilized by PyS2 for internalization. PyS2-488-stained PAO1 cells examined by confocal microscopy and super resolution structured illumination microscopy (SIM) revealed that the staining was localized to the outer envelope of PAO1 (Figure 5A) as expected. However, these observations did not discriminate between the outer membrane and periplasmic localization of PyS2-488.

**Figure 5.**
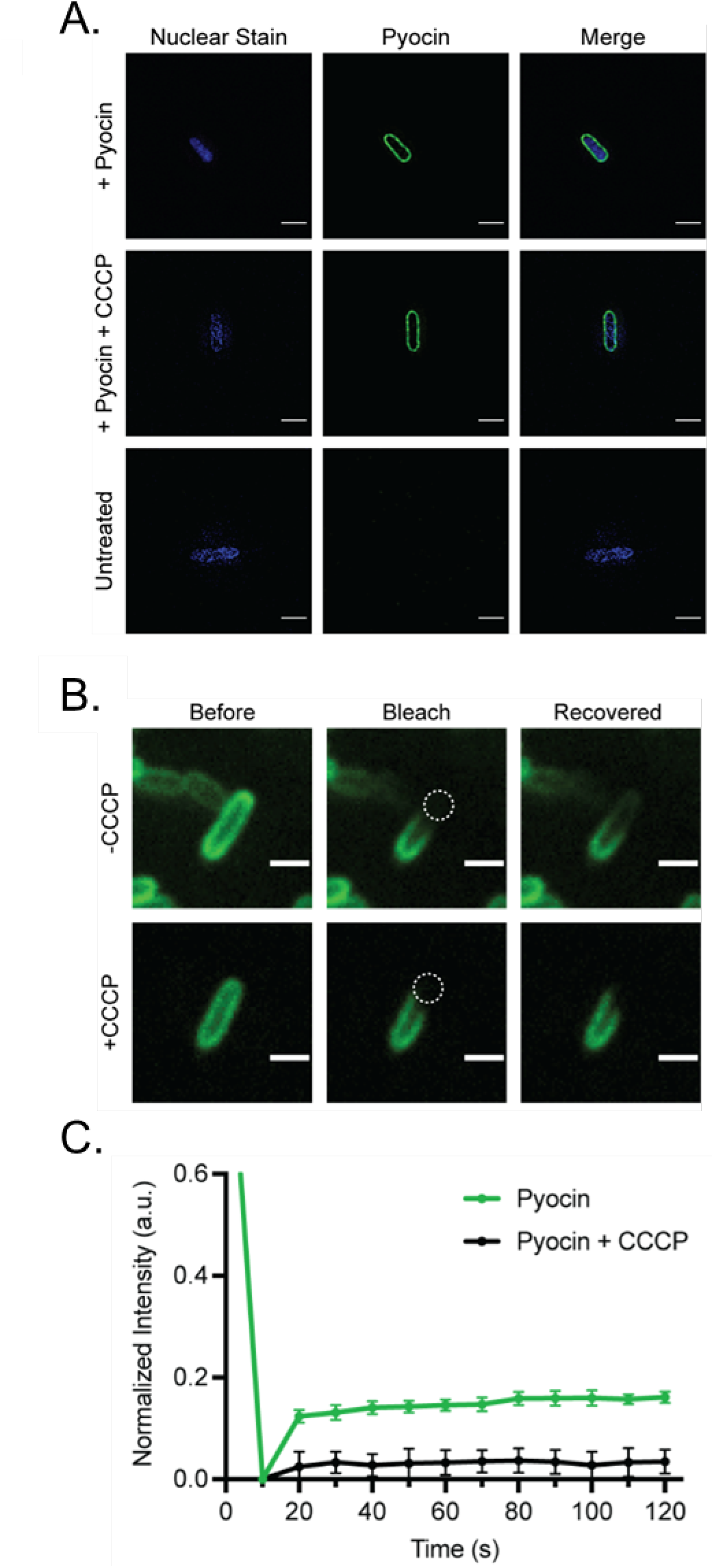
PyS2-488 binds *P. aeruginosa* and achieves periplasmic localization. PAO1 and PAO1 pretreated with 100 µM CCCP were stained with 1 µM PyS2-488 (see Supporting Information SI 13.2.) and examined with fluorescence microscopy techniques. **A**. Super resolution structured illumination microscopy (SIM) confirming periplasmic and outer membrane localization of the PyS2-488 (Scale bar = 3 µm, Excitation: 488 nm, Emission: 528 nm). **B**. Confocal fluorescence microscopy demonstrated localization of PyS2-488 to the outer membrane and periplasm and fluorescence recovery after photobleaching (FRAP) (Scale bar = 3 µm, Excitation: 488 nm, Emission: 525/40 nm). In both microscopy experiments SYTO 40 Blue was used as a nonspecific bacterial stain to localize and focus on the bacteria (Excitation: 405 nm, Emission: 435 nm for SIM and 447/60 nm for confocal). **C**. FRAP analysis illustrates PyS2-488 achieves periplasmic localization as a result of CCCP-dependent recovery.

To gain further insight into subcellular localization of PyS2-488, fluorescence recovery after photobleaching (FRAP) was used to determine if this staining was a result of binding at the outer membrane to FpvA or within the periplasm after internalization. If internalized to the periplasm, PyS2-488 would diffuse and recovery from FRAP would be observed. In contrast, the diffusion of outer membrane proteins is limited, therefore, FRAP recovery would not be observed if the fluorescently labeled protein is bound to FpvA on the outer membrane. PAO1 stained with PyS2-488 demonstrated partial rapid recovery, indicative of localization within the periplasm and ability to diffuse. In contrast, PAO1 cells pretreated with CCCP did not recover (Figure 5B), likely because most PyS2-488 was bound to the diffusion-limited outer membrane protein FpvA. Thus, we conclude that synthetic PyS2-488 was able to internalize into the *P. aeruginosa* periplasm. A large fraction of the fluorescence signal was bleached and did not recover relative to the original signal, suggesting that the majority of the PyS2-488 may remain bound to the FpvA receptor or undergoing transport. This fraction and recovery level is similar to those observed for recombinant PyS2^NTD^.^45^

## Conclusion

We report the single-shot solid-phase peptide synthesis of a 217 residue-long protein. The PyS2^NTD^ was manufactured in under 10 hours with an academic laboratory-scale automated fast-flow synthesizer and folded with and without fluorophore conjugation. The chemical synthesis of PyS2^NTD^ proceeded rapidly with high fidelity to yield a protein of unprecedented sequence length, further demonstrating the capabilities of automated flow synthesis technology.

The synthetic PyS2^NTD^ had similar biophysical and functional characteristics to those reported for the recombinant protein. Specifically, the CD signature and melting temperature closely matched the reported literature value for recombinant protein (T_m_ = 49 ± 2 °C observed, 50 ºC expected). Moreover, fluorophore-labeled PyS2-488 showed species- and FpvA receptor-specific binding and internalization. PyS2-488 staining supported the binding of PyS2-488 specifically to the FpvA receptor, evidenced by the limited staining of PAO1 Δ*fpvA, E. coli*, and PAO1 when grown with iron supplementation compared to the strong staining seen from PAO1 grown in iron-poor media. After FpvA-binding, synthetic PyS2-488 was localized in the periplasm, which required FpvA-coupled transport. Evidence for this transport was the transport inhibition by proton motive force inhibitor CCCP and its FRAP recovery in comparison to untreated PAO1.

Access to this class of pyocin proteins enables the exploration and potential development of non-traditional species-specific antimicrobials against *P. aeruginosa* infections. The low permeability of the outer membrane of *P. aeruginosa* is a major roadblock in developing new antibiotics. Although there are environment-dependent (ferric iron) effects of the internalization efficiency of PyS2^NTD^, the expression level of the ferripyoverdine receptor is related to the virulence of *P. aeruginosa*. At a minimum, pyocin-based antimicrobials stand to add selection pressure against the ferripyoverdine receptor, like inhibitors that have improved survival of the host *in vivo*.^57^

## Methods

See Supporting Information for all methods and supporting data including synthesis, cleavage, purification, folding, and conjugation conditions, as well as methods and supporting results for biophysical and biological characterization.

## Acknowledgements

This research was supported by a grant from the National Institute of Allergy and Infectious Diseases (U19 AI142780). A.S. gratefully acknowledges support from the National Science Foundation Graduate Research Fellowship under Grant no. 1122374. J.S.B. acknowledges support from the Pharmaceutical Research and Manufacturers of America (PhRMA) through the Postdoctoral Fellowship in Drug Discovery. V.M.M. acknowledges support from the Natural Sciences and Engineering Research Council of Canada (NSERC) through the PGSD Fellowship. The authors also thank Alex Callahan for helpful discussions on protein folding and the Koch Institute’s Robert A. Swanson (1969) Biotechnology Center for technical support (NCI Cancer Center Support Grant P30 CA14051), specifically The Microscopy Core Facility and Dr. Jeffrey R. Kuhn. Also, we thank the W.M. Keck imaging facility at the Whitehead Institute for access to the ANDOR FRAPPA confocal microscope.

