## Supporting Information for "Rapid Single-Shot Synthesis of the 217 Amino Acid-Long N-Terminal Domain of Pyocin S2"

**Table of Contents**

Rapid Single-Shot Synthesis of the 217 Amino Acid-Long N-Terminal Domain of Pyocin S2 ... 1
SI 1.0. Materials ... 3
SI 2.0. Analysis methods with Liquid-Chromatography Mass Spectrometry (LC-MS) ... 4
SI 3.0. PyS2$^{NTD}$ characteristics ... 6
SI 4.0. PyS2$^{NTD}$ Synthesis and Cleavage ... 7
Reagent storage ... 7
Pumping, mixing, and heat exchange ... 7
Synthesis sequence. ... 8
Cleavage. ... 9
Yield calculation ... 10
SI 5.0. Characterization of crude PyS2$^{NTD}$ ... 12
SI 5.1. Analytical HPLC of crude PyS2$^{NTD}$ ... 13
SI 5.2. LCMS of Crude PyS2$^{NTD}$ ... 15
SI 6.0. HPLC Purification of PyS2$^{NTD}$ ... 16
SI 6.1. HPLC purification of crude PyS2$^{NTD}$ ... 18
SI 6.2. LC-MS analysis of fractions from RP-HPLC purification of PyS2$^{NTD}$ ... 20
SI 7.0 Characterization of HPLC-purified PyS2$^{NTD}$ ... 27
SI 7.1. Analytical HPLC of HPLC-purified PyS2$^{NTD}$ ... 28
LCMS of HPLC-purified PyS2$^{NTD}$ ... 30
SI 8.0. PyS2$^{NTD}$ Folding ... 31
SI 8.1. SEC chromatogram of folding of PyS2$^{NTD}$ ... 33
SI 9.0. Characterization of folded PyS2$^{NTD}$ ... 35
SI 9.1. Analytical HPLC of folded PyS2$^{NTD}$ ... 36
SI 9.2. LC-MS of folded PyS2$^{NTD}$ ... 38
SI 10.0. Circular dichroism measurements ... 39
SI 11.0. Fluorophore conjugation of PyS2$^{NTD}$ ... 40
Reaction conditions. ... 40
SI 11.1. Monitoring reaction by LC-MS ... 42
SI 11.2. SEC chromatogram of purification/folding of PyS2-488 ... 43
SI 12.0. Characterization of PyS2-488 ... 45
SI 12.1. Analytical HPLC analysis of PyS2-488 ... 46
SI 12.2. LC-MS of PyS2-488 ... 48
SI 13.0. Biological Characterization of PyS2$^{NTD}$ ... 49
SI 13.1. Bacterial Strains and Construction ... 49
SI 13.2. Staining of *Pseudomonas aeruginosa* with PyS2$^{NTD}$-fluorophore conjugate ... 49
SI 13.3. Flow cytometry ... 50
SI 13.4. Confocal microscopy and fluorescence recovery after photobleaching (FRAP) ... 50
SI 14.0. References ... 56

### SI 1.0. Materials

All reagents were purchased and used as received unless stated otherwise. Fmoc-protected amino acids Fmoc-L-Ala-OHxH2O, Fmoc-L-Arg(Pbf)-OH; Fmoc-L-Asn(Trt)-OH; Fmoc-L-Gln(Trt)-OH; Fmoc-L-Ile-OH; Fmoc-L-Leu-OH; Fmoc-L-Lys(Boc)-OH; Fmoc-L-Pro-OH; Fmoc-L-Ser(*t*-Bu)-OH; Fmoc-L-Tyr(*t*-Bu)-OH, were purchased from Sigma Millipore (Novabiochem) and lyophilized for at least 24 hours prior to use. Fmoc-protected amino acids Fmoc-L-Asp-(O*t*-Bu)-OH; Fmoc-L-Cys(Trt)-OH; Fmoc-L-Glu(O*t*-Bu)-OH; Fmoc-Gly-OH; Fmoc-L-Phe-OH; Fmoc-L-Thr(*t*-Bu)-OH; Fmoc-L-Val-OH), were purchased from Sigma Millipore (Novabiochem) and used as received. Fmoc-L-Nle-OH was purchased from Chem-Impex and lyophilized for at least 24 hours prior to use. Fmoc-L-His(Boc)-OH was purchased from Advanced ChemTech and used as received. O-(7-azabenzotriazol-1-yl)-*N*,*N*,*N*',*N*'-tetramethyluronium hexafluorophosphate (HATU, ≥97.0%) and (7-azabenzotriazol-1-yloxy)tripyrrolidinophosphonium hexa-fluorophosphate (PyAOP, ≥97.0%) were purchased from P3 Biosystems. Biosynthesis OmniSolv® grade *N*,*N*-dimethylformamide (DMF) was purchased from EMD Millipore (DX1732-1) and incubated with 2 packs of AldraAmine trapping agents (for 1000–4000 mL volume, Sigma-Aldrich, catalog number Z511706) for 48 hours prior to use. Diisopropylethylamine (DIEA; 99.5%, biotech grade, catalog number 387649) and piperidine (ACS reagent, ≥99.0%) were purchased from Sigma-Aldrich. Formic acid (FA, 97%) was purchased from Beantown Chemical, corp. Trifluoroacetic acid (HPLC grade, ≥99.0%), Phenol (ACS reagent, 99.0-100.5%), Thioanisole (ReagentPlus®, ≥99%), Diethyl ether (anhydrous, ACS reagent, ≥99.0%), acetonitrile (HPLC grade, ≥99.9%), Omnisolv® acetonitrile (LC-MS grade, AX0156-1), Omnisolv® water (LC-MS grade, WX0001-1) and dimethyl sulfoxide (DMSO, molecular biology grade, D8418) were purchased from Sigma-Aldrich. Formic acid Optima

LC/MS (A117) was purchased from Fisher Chemical. H-Rink Amide-Chemmatrix® LL (0.17 mmol/g) resin was purchased from PCAS Biomatrix. Water was deionized using a Milli-Q Reference water purification system (Millipore). Nylon 0.22 μm syringe filters were TISCH brand SPEC17984.

**SI 2.0. Analysis methods with Liquid-Chromatography Mass Spectrometry (LC-MS)**

LC-MS analysis with methods A and B were acquired using an Agilent G6545A MS Q-TOF mass spectrometer with Dual AJS ESI ion source in extended dynamic mode in mass range 100-3000 *m/z* with scan rate of 1.00 spectra/sec. The following instrument parameters were used: gas temperature 350 °C, gas flow 11 L/min, nebulizer pressure 60 psig, sheath gas temperature 350 °C, sheath gas flow 11 L/min. The following scan source parameters were used: VCap: 3500, nozzle voltage 1000 V, fragmentor 175, skimmer1 65, OctopoleRFPeak 750. The following LC parameters were used: solvent system 0.1% formic acid in $H_2O$ (solvent A) and 0.1% formic acid in $CH_3CN$ (solvent B), flow rate 0.8 mL/minute, column Zorbax 300SB C3 2.1 × 150 mm, 5 μm), column temperature 40 °C.

**Method A (6545-15min)** gradient: 0-4 minutes 1% B, 4-11 minutes 1-91% B, 11-15 minutes 91-95% B. LC stream directed to the MS in 4-11 minutes.

**Method B (6545-15min)** gradient: 0-1 minutes 1% B, 1-7 minutes 1-91% B, 7-9 minutes 91-95% B. LC stream directed to the MS in 2-7 minutes.

Data were processed using Agilent MassHunter BioConfirm Software 10.0. Deconvoluted masses of proteins were obtained using a maximum entropy algorithm. For deconvolution a range with lower limit of 10,000 Da and upper limit of 60,000 Da was used. A limited *m/z* range from

600-3,000 was used with a baseline subtraction factor of 3 and a mass step of 1. The 15,000-30,000 range has been plotted in most figures. Y-axis in all chromatograms shown in Supporting Figures represents total ion current (TIC); mass spectrum insets correspond to the deconvolution of the entirety of the protein containing peaks.

LC-MS analysis with method C was acquired using an Agilent G6550A MS Q-TOF mass spectrometer with Dual AJS ESI ion source in extended dynamic mode in mass range 100-3000 *m/z* with scan rate of 1.00 spectra/sec. The following instrument parameters were used: gas temperature 200 °C, gas flow 14 L/min, nebulizer pressure 55 psig, sheath gas temperature 350 °C, sheath gas flow 11 L/min. The following scan source parameters were used: VCap: 5500, nozzle voltage 2000 V, fragmentor 380, skimmer1 65, OctopoleRFPeak 750. The following LC parameters were used: solvent system 0.1% formic acid in $H_2O$ (solvent A) and 0.1% formic acid in $CH_3CN$ (solvent B), flow rate 0.8 mL/minute, column Zorbax 300SB C3 2.1 × 150 mm, 5 µm), column temperature 40 °C.

**Method C (6550-15min)** gradient: 0-2 minutes 1% B, 2-12 minutes 1-61% B, 12-12.1 minutes 61-90% B. LC stream directed to the MS in 2-12 minutes.

Data were processed using Agilent MassHunter Qualitative Analysis Software B.06.00. Deconvoluted masses of proteins were obtained using the same method as A and B.

**SI 3.0. PyS2$^{NTD}$ characteristics**

**Sequence:**

$NH_2$-

MAVNDYEPGSMVITHVQGGGRDIIQYIPARSSYGTPPFVPPGPSPYVGTGMQEYRKLRST

LDKSHSELKKNLKNETLKEVDELKSEAGLPGKAVSANDIRDEKSIVDALMDAKAKSLK

AIEDRPANLYTASDFPQKSESMYQSQLLASRKFYGEFLDRHMSELAKAYSADIYKAQIAI

LKQTSQELENKARSLEAEAQRAAAEVEADYK**GGSCG**-$CONH_2$

*M = Norleucine (Nle)

**Bolded sequence was appended to the native PyS2$^{NTD}$ sequence as a soluble spacer and bioconjugation handle.

**Calculated molecular weight of PyS2$^{NTD}$:** 23435 Da

**Calculated molecular weight of trifluoroacetate salt of PyS2$^{NTD}$** (+33 TFA counter ion molecules from protonation of 19 lysine residues, 3 histidine residues, 10 arginine residues and the N-terminal amine): 27198 Da

### SI 4.0. PyS2$^{NTD}$ Synthesis and Cleavage

The PyS2$^{NTD}$ sequence synthesized on pre-swollen 100 mg of LL ChemMatrix Rink Amide resin (loading 0.17 mmol/g) using the published optimized protocol for automated flow peptide synthesis (AFPS) as described previously.[1]

**Reagent storage.** The reagent storage system for amino acids, coupling agents, and deprotection solution used Chemglass three-neck 200- or 500-ml spinner flasks. Reagent withdraw lines were outfitted with a 20 μm polypropylene filter (Valco part # JR-32178) to prevent clogging of pumps, check valves, and lines from any reagent crystallization or impurities. The amino acids were selected using a VICI Valco 10-position auxiliary valve fed into the main valve connected to the amino acid pump. Bottles containing HATU, other coupling agents, 40% piperidine and DMF fed into a separate 10-position valve, the coupling agent valve. This valve was connected to the coupling agent pump. DIEA fed directly into the DIEA Pump. The tubing lengths are such that the volume between the coupling agent pump and the coupling agent valve was equal to the volumes between the amino acid pump and each auxiliary valve. This is crucial to avoid truncations of the peptide chain, ensuring that, at the start of each coupling cycle, amino acid arrived at the mixer at the same time as activating agent.

**Pumping, mixing, and heat exchange.** Amino acids (0.4 M) in amine-free DMF and 0.38 M activators in amine-free DMF were combined using autonomously controlled HPLC pumps (Varian Prostar 210, run at 80% of maximum speed giving 40 mL/min from a 50 mL/min pump head, except DIEA which gives 4 mL/min with a 5 mL pump head) into check valves (IDEX part # CV-3320) connected to a mixing tee (IDEX part # P-722) using PEEK tubing (1/16″ OD, 0.020″ ID). After the mixing tee, a differential pressure sensor was installed to monitor the heat exchanger and reactor pressure (DJ Instrument, Through-bore titanium pressure sensors, part # DF2-01-TI-500-5V-41″). Then, PEEK tubing was coiled to form a high Dean number (>3,000) static mixer to facilitate reagent mixing. The PEEK tubing delivered the mixed solution to a VICI Valco 6-position column selector valve (Vici part# ACST6UW-EUTA) to be directed to stainless steel heat exchangers, consisting of stainless-steel tubes wrapped around a steel core heated with a heated with two resistive cartridge heaters (Omega part # CSS-10250/120V). Two heat exchangers were used and operated at 90 °C, 10' heating loop (default coupling) or 60 °C, 5' heating loop (His and Cys only). The outflow of the heat exchangers was directed to a 90 °C steel block reactor inlet, which inserts into a 6 ml HSW Norm-Ject syringe (Torviq), with a pre-inserted coarse polyethylene frit and filter paper to retain the ChemMatrix resin under flow. The reactor is jacketed by a steel block heated to 85 °C with cartridge heaters actuated by a robotic arm. After the reactor, UV monitoring at 312 nm was accomplished by using a Varian Prostar 230 UV-visible detector before passing to waste.

**Synthesis sequence.** Amine-free DMF was passed over the resin to wash the resin before coupling, after coupling to push through excess reagents, and after deprotection (40 strokes, ~25 mL). Coupling was performed with HATU (single-coupling, 8 strokes, ~5 mL) except S&A with HATU (double-coupling, 21 strokes, ~10 mL) and C, H, N, Q, R, V, T with PyAOP (double-coupling, 21 strokes, ~10 mL). Wash was repeated. Deprotection was completed with 20% piperidine in amine-free DMF with 2% formic acid (2 pumps, 40 mL/min), followed by a wash sequence. Amino acids are coupled with this recipe in iterative sequence until complete. A final double wash of amine-free DMF was completed after the last deprotection step. LabView was used to program the sequence of coupling, deprotection, and washing steps.

**Cleavage.** After synthesis, the peptidyl resin was washed with DMF (3 x 5 mL) and with dichloromethane (3 x 5 mL), dried in a vacuum chamber, and weighed. The resin was then transferred into a 20 mL fritted syringe. 15 mL of reagent K cleavage solution (82.5% TFA, 5% water, 5% phenol, 5% thioanisole, 2.5% EDT) was added the syringe to cover the resin. The syringe was then capped and kept on a nutating mixer at room temperature for 4 h. Next, the mixture was ejected out of the syringe, separating the resin from the solution. The resin was washed with another 5 mL of reagent K cleavage solution and the solutions were combined, yielding total approximately 20 mL of cleavage mixture. The resin was discarded. Equal volume of the mixture was added to two conical polypropylene (falcon) tubes. Cold diethyl ether (45 mL) chilled by keeping on dry ice, was added to each tube containing the cleavage mixture. The resulting white precipitate was collected by centrifugation and triturated twice more with cold diethyl ether (45 mL). The supernatant was discarded. To the precipitate, 5 mL of 50% acetonitrile in water with 0.1% TFA was added. The mixture was vortexed briefly (<1 min), but no further efforts to fully dissolve the material were made. The mixture was flash frozen with liquid nitrogen and lyophilized until dry, typically for 12-24 hours, to yield the crude $PyS2^{NTD}$.

**Yield calculation.** For a typical synthesis, the mass of peptidyl resin was 587 mg and the mass of crude PyS2$^{NTD}$ was 197 mg, leading to 43% yield. The synthesis yield was calculated based on the theoretical yield from the resin loading, versus the amount isolated following cleavage and lyophilization.

$$synthesis\ yield\ (\%) = \frac{isolated\ amount\ of\ crude\ material\ (mg)}{amount\ resin\ used\ (g) \times\ resin\ loading\ \left(\frac{mmol}{g}\right) \times molecular\ weight\ (\frac{g}{mol})} \times 100$$

Note that the molecular weight used in this formula accounts for 33 trifluoroacetate counter ions; See section on "PyS2$^{NTD}$ characteristics".

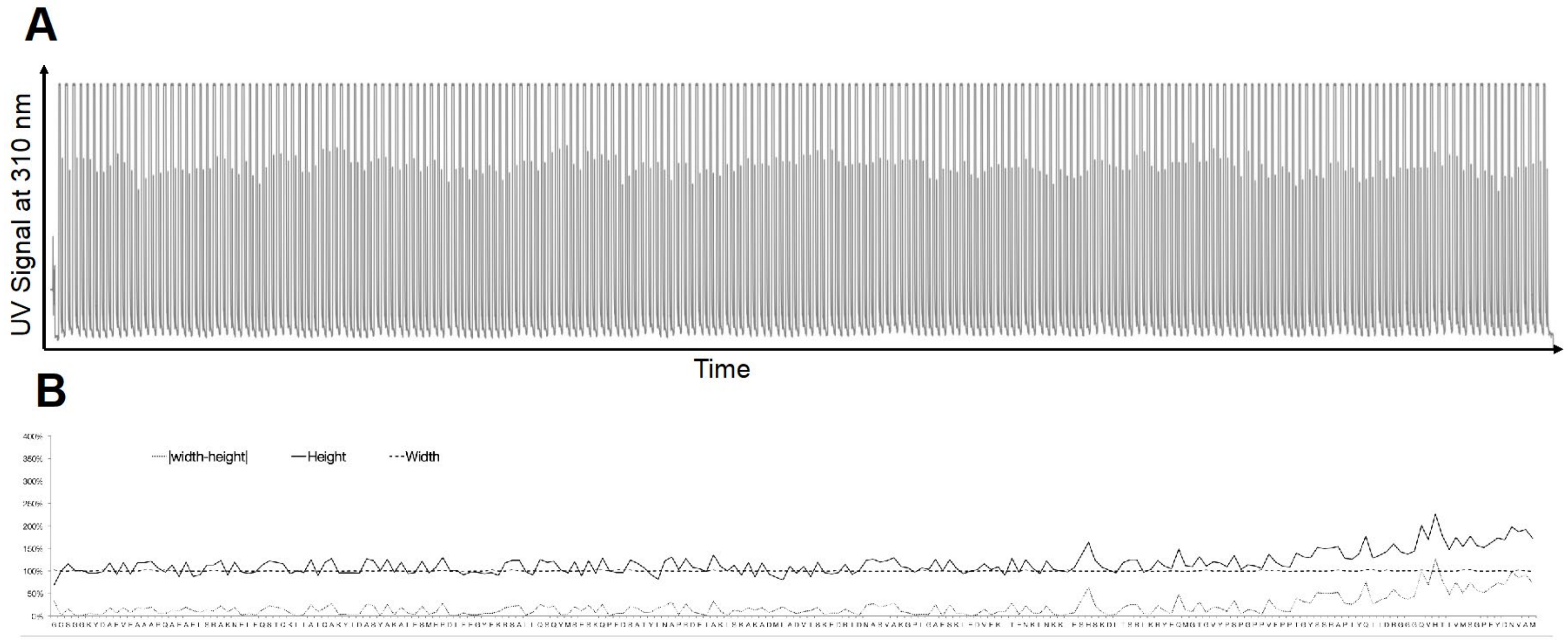


**Figure S1. A.** The synthesis trace of PyS2$^{NTD}$ is depicted showing no major changes in height of Fmoc deprotection peaks throughout the synthesis. **B.** Analysis of deprotection peaks shows minor changes in height and width of deprotection peaks, suggesting no major aggregation events occurred during synthesis of the sequence.

### SI 5.0. Characterization of crude PyS2$^{NTD}$

Crude PyS2$^{NTD}$ was characterized by HPLC and LC-MS.

For assessment of purity by HPLC, crude PyS2$^{NTD}$ was dissolved in a freshly prepared solution of 1:1 acetonitrile:water with 0.1% TFA and 500 mM DTT, to yield a solution of PyS2$^{NTD}$ (at minimum 50 µL) at approximately 1.0 mg/mL. The solution was filtered and analyzed on an analytical Agilent Zorbax 300SB-C3 column (2.1 mm × 150 mm, 5-µm particle size) on Agilent Technologies 1200 Series instrument, which was computer-controlled through Agilent ChemStation software. A linear gradient of acetonitrile with a 0.08% TFA additive (solvent B) in water with a 0.1% TFA additive (solvent A) was used. Gradients of 5% B per minute ramped up to 65% over 30 min at a flow rate of 0.4 mL/min. Wash and recalibration of the column sequence included ramp up of gradient to 100% B at 31 min, hold at 100% B until 36 min, ramp down to 5% B at 37 min, and hold at 5% B until 43 min. The total method time was 43 min.

For LC-MS method A was used as described in "Analysis method with Liquid-Chromatography Mass Spectrometry (LC-MS)" section.

**SI 5.1. Analytical HPLC of crude PyS2$^{NTD}$**

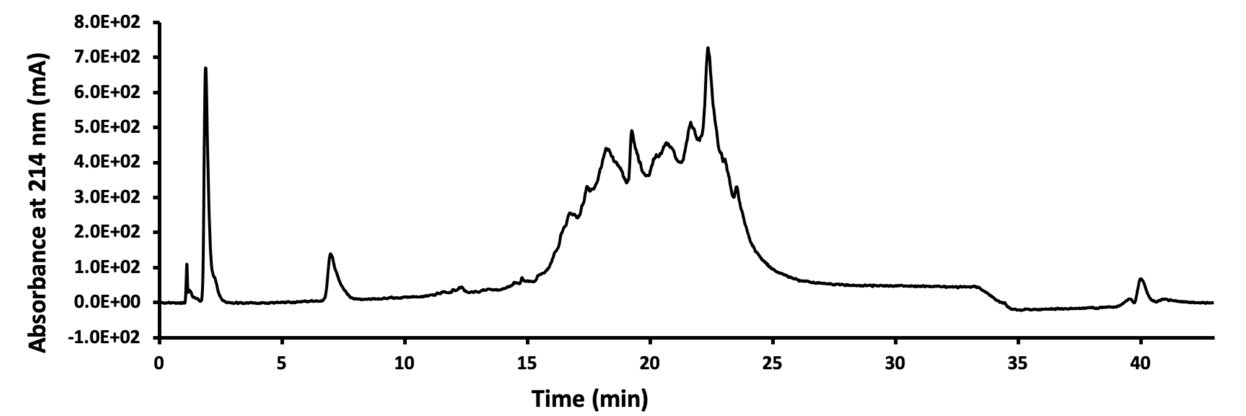


**Figure S2.** HPLC chromatogram of the crude PyS2$^{NTD}$ displaying the absorbance at 214 nm.

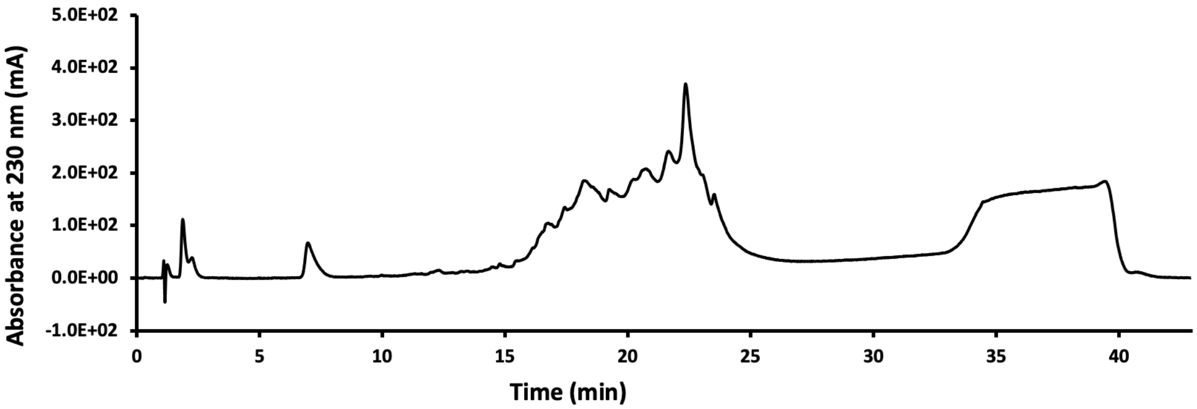


**Figure S3.** HPLC chromatogram of the crude PyS2$^{NTD}$ displaying the absorbance at 230 nm.

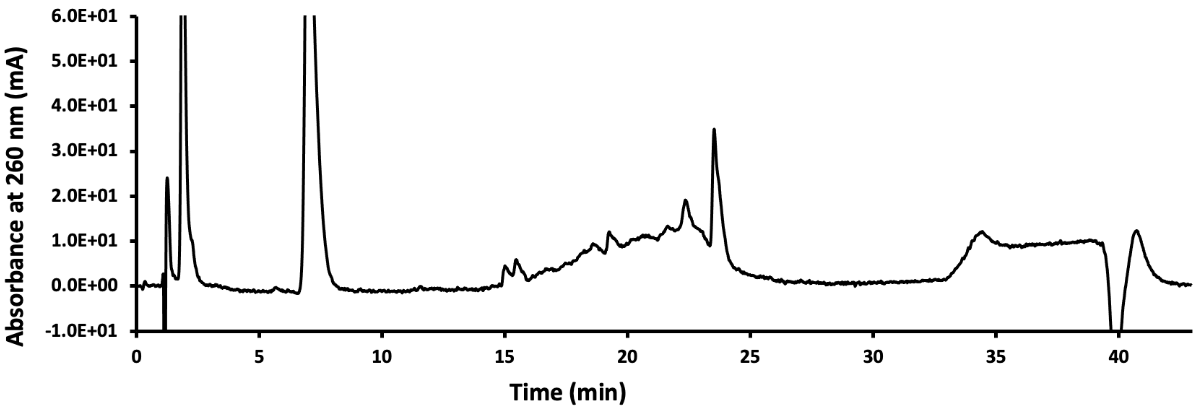


**Figure S4.** HPLC chromatogram of the crude PyS2$^{NTD}$ displaying the absorbance at 260 nm.

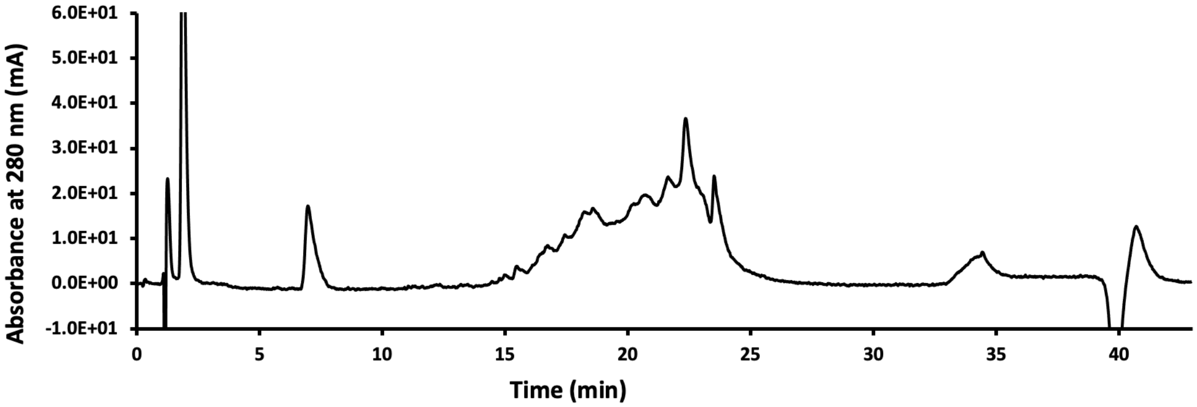


**Figure S5.** HPLC chromatogram of the crude PyS2$^{NTD}$ displaying the absorbance at 280 nm.

### SI 5.2. LCMS of Crude PyS2$^{NTD}$

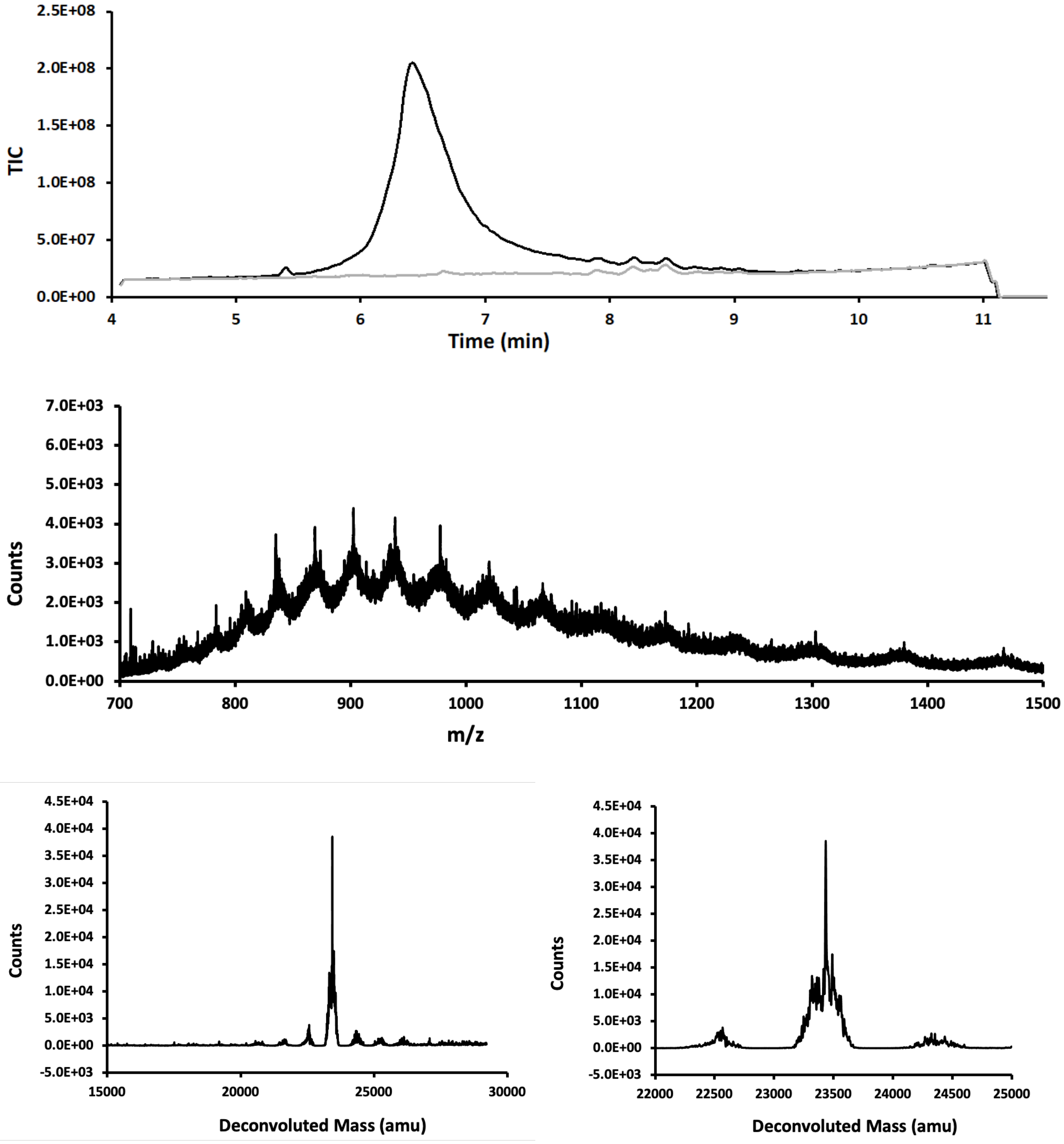


**Figure S6.** LC-MS analysis of crude PyS2$^{NTD}$. Top panel is the total ion count versus time. In grey spectrum of a blank run is overlayed. The mass spectrum in the middle panel is obtained from integration over the entire peak and is presented in m/z (in Da) versus time. Bottom panels display deconvoluted mass spectra. Calc.:23435 Da, Obs. 23423 Da.

### SI 6.0. HPLC Purification of PyS2$^{NTD}$

The lyophilized crude sample of PyS2$^{NTD}$ was weighed and portioned equally into two 50 mL conical tubes. Each portion was purified separately, simply to avoid complete sample loss due to potential instrument errors. Prior to HPLC purification, to the crude sample of PyS2$^{NTD}$, 10 mL of solution of 6 M guanidinium chloride, 0.1 M dithiothreitol (DTT), in 50 mM sodium phosphate pH 7.5 was added, the mixture was vortexed briefly and then filtered and subjected to RP-HPLC purification. For this purification, the Agilent Zorbax 300SB-C18 PrepHT (21.2 × 250 mm, 7 µm) heated at 60 °C, used at 20.0 mL/min with the following gradient: isocratic 5% B from 0–5 min; linear gradient from 5–35% B from 5–12 min; isocratic 35% B from 12–15 min; linear gradient from 35–45% B from 15–45 min; linear gradient from 45–65% B from 45–50 min; isocratic 65% B from 50–55 min. During 15–45 min, 15-second aliquots were collected and analyzed by LC-MS. Fractions showing high purity charge state series were combined and lyophilized (Figure S12).

Using this procedure, purification of 182 mg of crude PyS2$^{NTD}$ (trifluoroacetate salt) led to isolation of 10.3 mg of HPLC-purified PyS2$^{NTD}$ (trifluoroacetate salt), amounting to 6% yield. Generally, a typical synthesis consistently led to 8-11 mg of HPLC-purified PyS2$^{NTD}$. The isolation yield was calculated based on the amount of HPLC-purified material isolated, versus the amount of crude material used.

$$isolation\ yield\ (\%) = \frac{HPLC - purified\ material\ (mg)}{crude\ material\ (mg)} \times 100$$

| Time (min) | Percent solvent B (MeCN + 0.1% TFA) (%) |
|---|---|
| 0 | 5 |
| 5 | 5 |
| 12 | 35 |
| 15 | 35 |
| 45 | 45 |
| 50 | 65 |
| 55 | 65 |

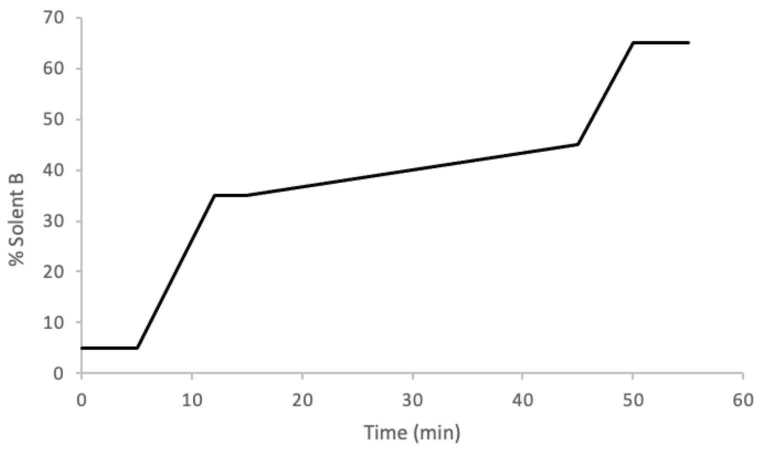


**Figure S7.** Gradient of solvent B (acetonitrile with 0.1% TFA) for RP-HPLC of crude $PyS2^{NTD}$.

#### SI 6.1. HPLC purification of crude PyS2$^{NTD}$

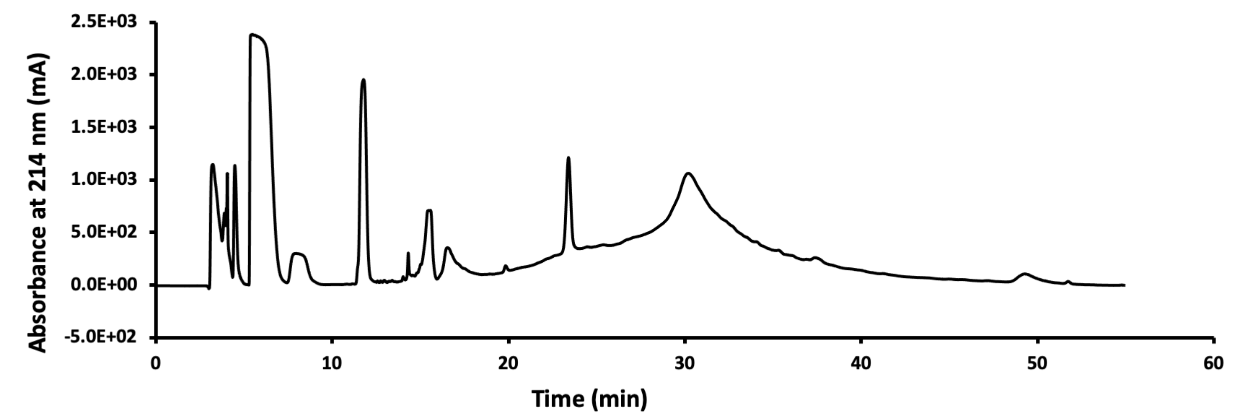


**Figure S8.** Chromatogram from purification of PyS2$^{NTD}$ with preparative reverse-phase high performance liquid chromatography (RP-HPLC) displaying the absorbance at 214 nm. Product elutes around the 30-minute mark. Fractions containing the product, as determined by LC-MS, were combined, and lyophilized to afford the HPLC-purified PyS2$^{NTD}$ as trifluoroacetate salt.

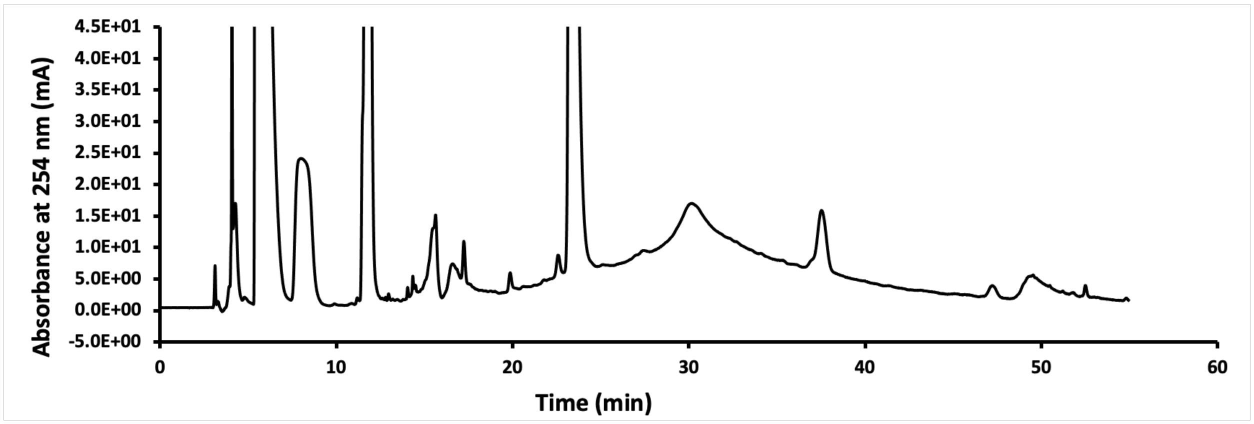


**Figure S9.** Chromatogram from purification of PyS2$^{NTD}$ with preparative reverse-phase high performance liquid chromatography (RP-HPLC) displaying the absorbance at 254 nm.

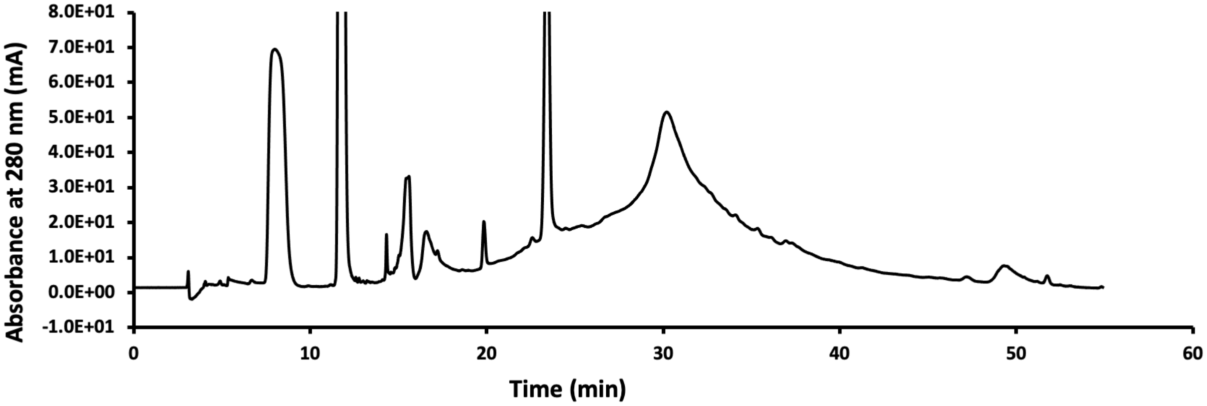


**Figure S10.** Chromatogram from purification of PyS2$^{NTD}$ with preparative reverse-phase high performance liquid chromatography (RP-HPLC) displaying the absorbance at 280 nm.

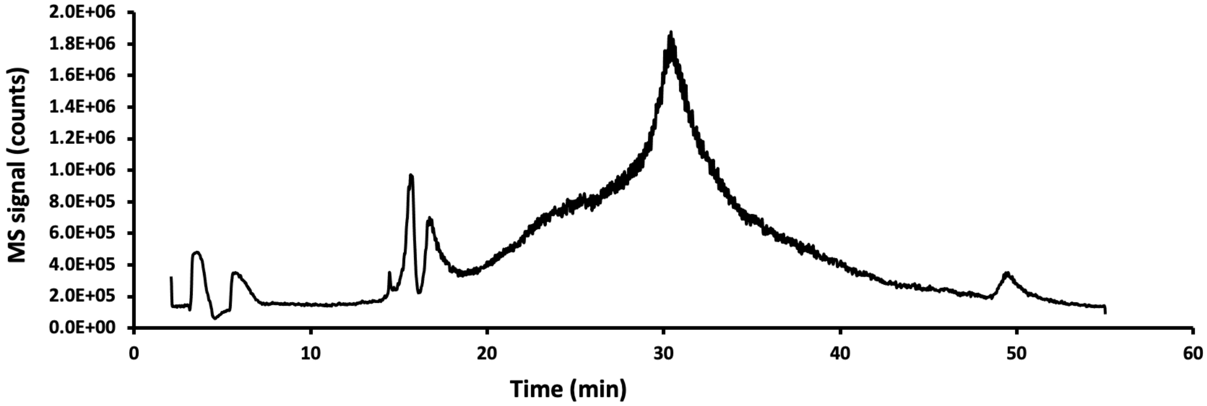


**Figure S11.** Chromatogram from purification of PyS2$^{NTD}$ with preparative reverse-phase high performance liquid chromatography (RP-HPLC) displaying the MS signal.

#### SI 6.2. LC-MS analysis of fractions from RP-HPLC purification of PyS2$^{NTD}$

Fractions from RP-HPLC purification were collected in 0.25 min time intervals. Guided by the 214 nm absorbance signal from elution of the product, fractions with high absorbance were analyzed by LC-MS. Specifically 30 μL of each fraction with suspected accumulated product was transferred to a vial for LC-MS analysis without further dilution. Typically, 1-2 μL of injection of the solution using method B resulted in sufficient signal for analysis (figures S13-17). The fractions were kept at 4 °C during LC-MS analysis, typically overnight but not over 24 hours. The fractions with highest level quality of MS spectrum and deconvoluted mass spectrum were combined, flash frozen and lyophilized. The product was thereafter stored in freezer at -80 °C.

Figures S13-17 depict a typical LC-MS analysis done on fractions. In this case, the two tailing fractions (fraction 64, 68) were of sufficient purity and thus discarded. Fractions 65, 66, 67 were combined to result in the final HPLC-purified PyS2$^{NTD}$. The overlay of the fractions collected and the 214 nm absorbance signal is depicted in figure S12.

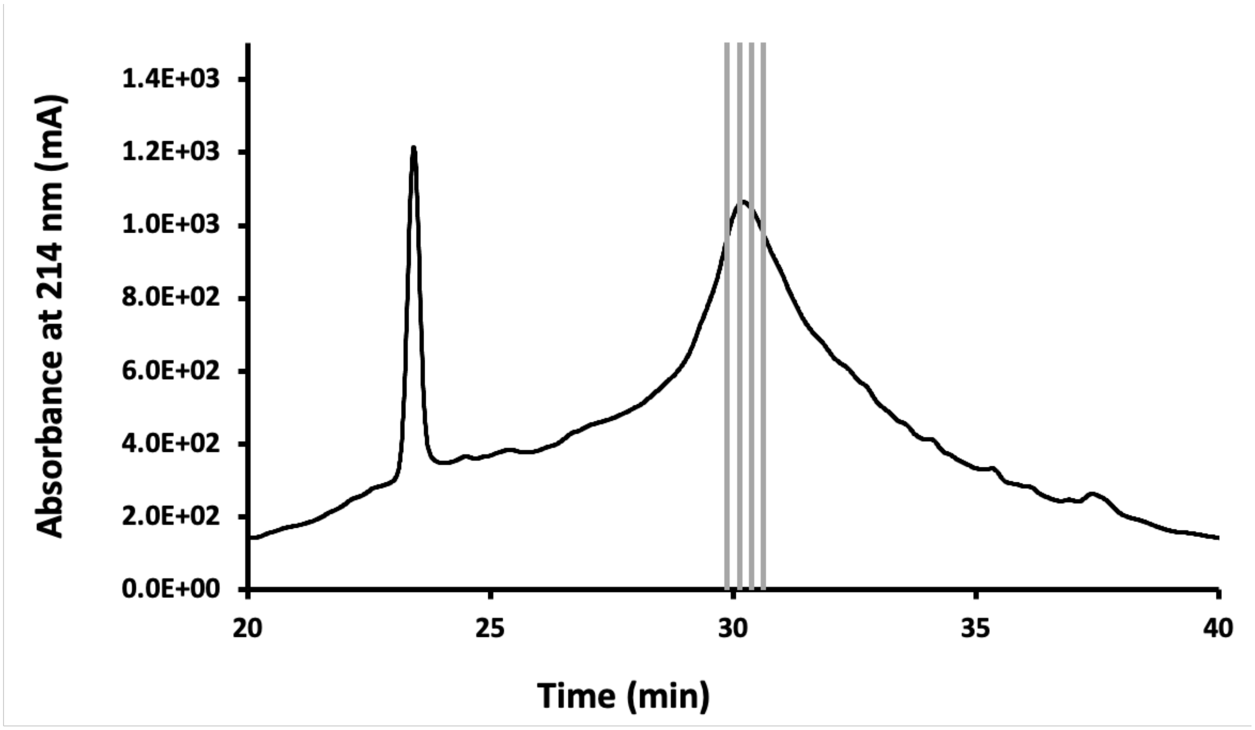


**Figure S12.** Chromatogram from purification of $PyS2^{NTD}$ with RP-HPLC displaying the absorbance at 214 nm in the 20-40 min range and the fractions that were combined to result in the final product. Fractions were analyzed by LC-MS to determine if they have sufficient purity (Figures S13-17).

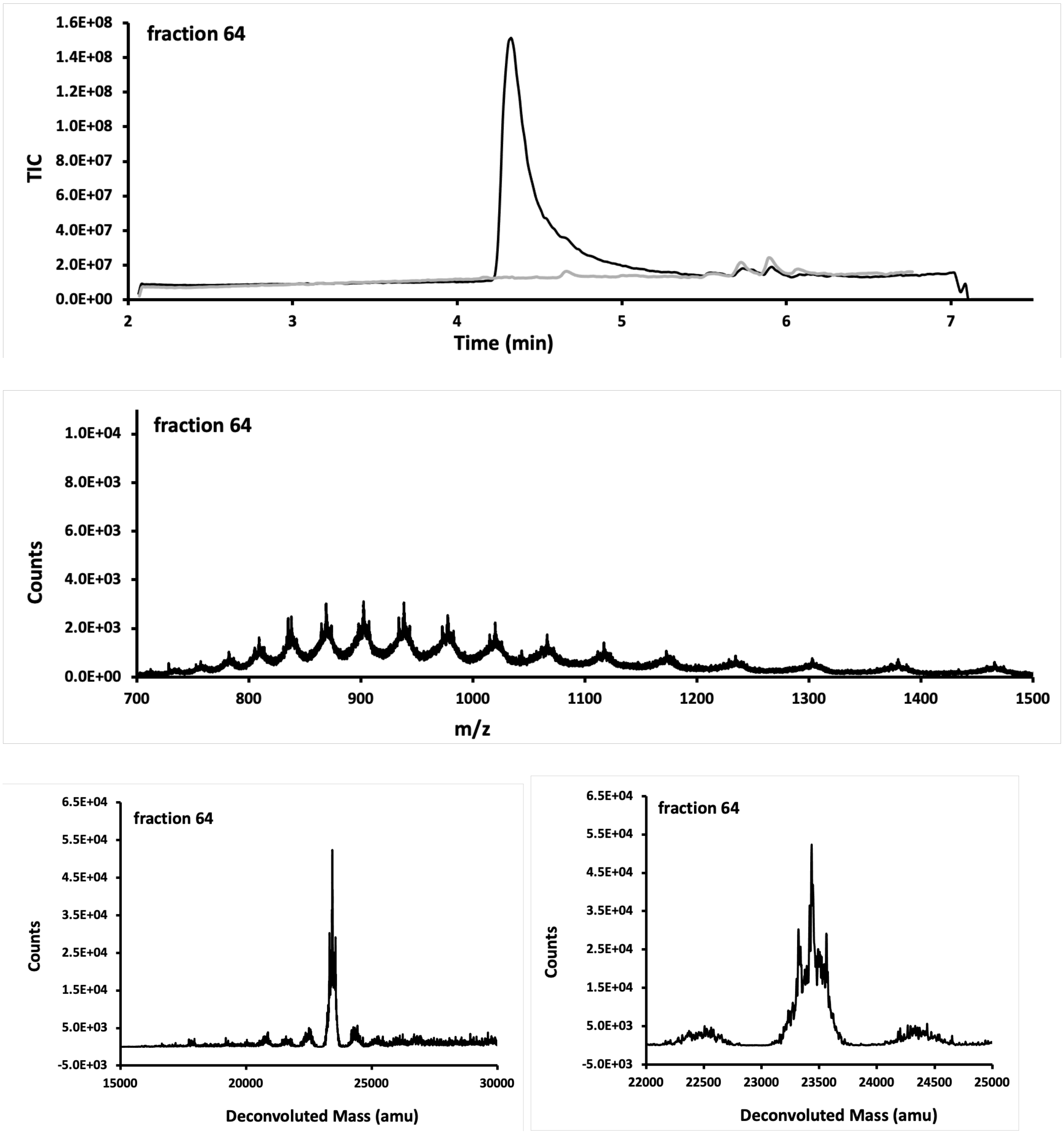


**Figure S13.** LC-MS analysis of fraction 64 of PyS2$^{NTD}$. Top panel is the total ion count versus time. In grey spectrum of a blank run is overlayed. The mass spectrum in the middle panel is obtained from integration over the entire peak and is presented in m/z (in Da) versus time. Bottom panels display deconvoluted mass spectra. This fraction did not exhibit enough purity and was discarded. Calc.:23435 Da, Obs. 23423 Da.

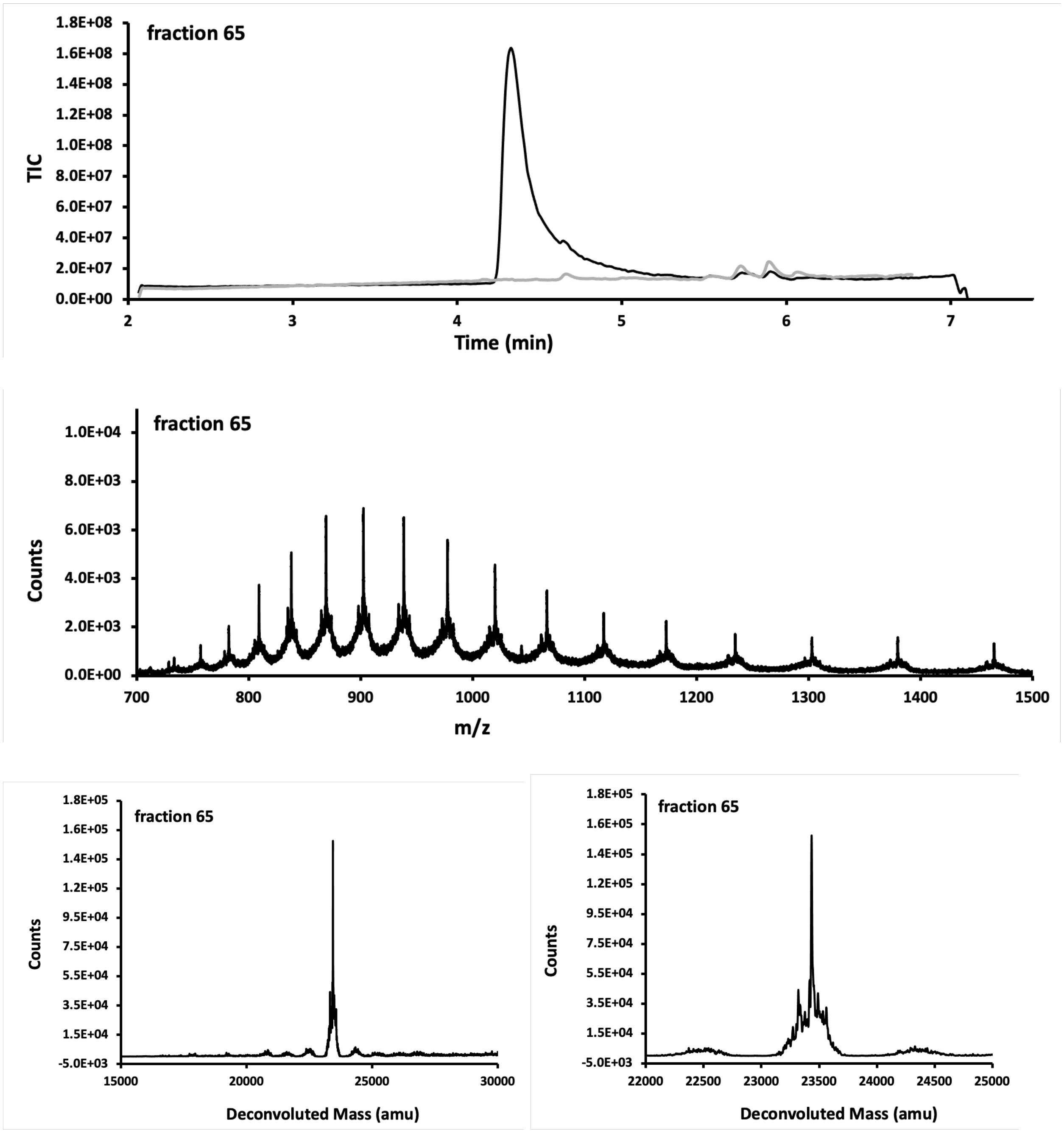


**Figure S14.** LC-MS analysis of fraction 65 of PyS2$^{NTD}$. Top panel is the total ion count versus time. In grey spectrum of a blank run is overlayed. The mass spectrum in the middle panel is obtained from integration over the entire peak and is presented in m/z (in Da) versus time. Bottom panels display deconvoluted mass spectra. This fraction combined to yield the final product. Calc.:23435 Da, Obs. 23423 Da.

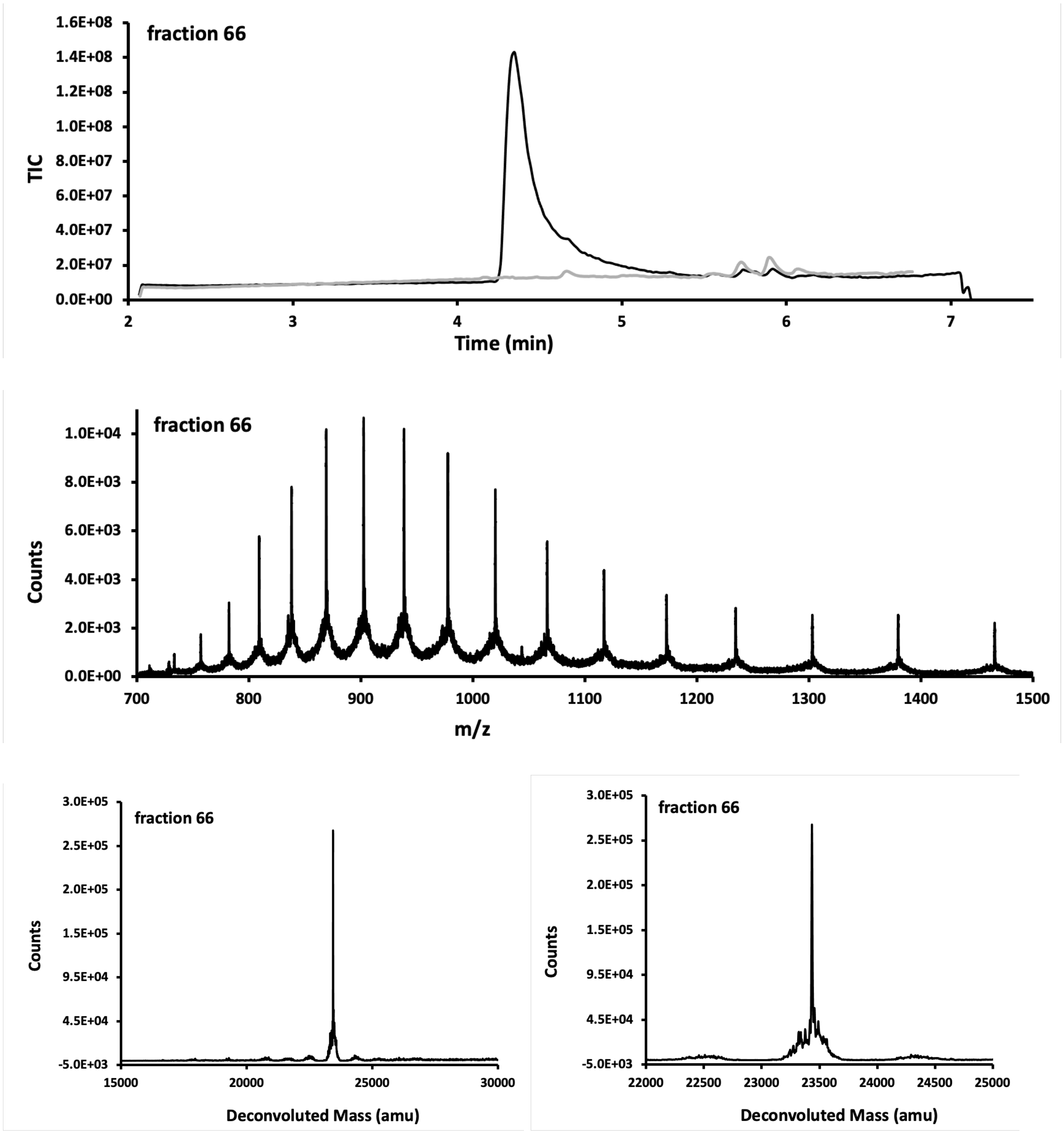


**Figure S15.** LC-MS analysis of fraction 66 of PyS2$^{NTD}$. Top panel is the total ion count versus time. In grey spectrum of a blank run is overlayed. The mass spectrum in the middle panel is obtained from integration over the entire peak and is presented in m/z (in Da) versus time. Bottom panels display deconvoluted mass spectra. This fraction combined to yield the final product. Calc.:23435 Da, Obs. 23423 Da.

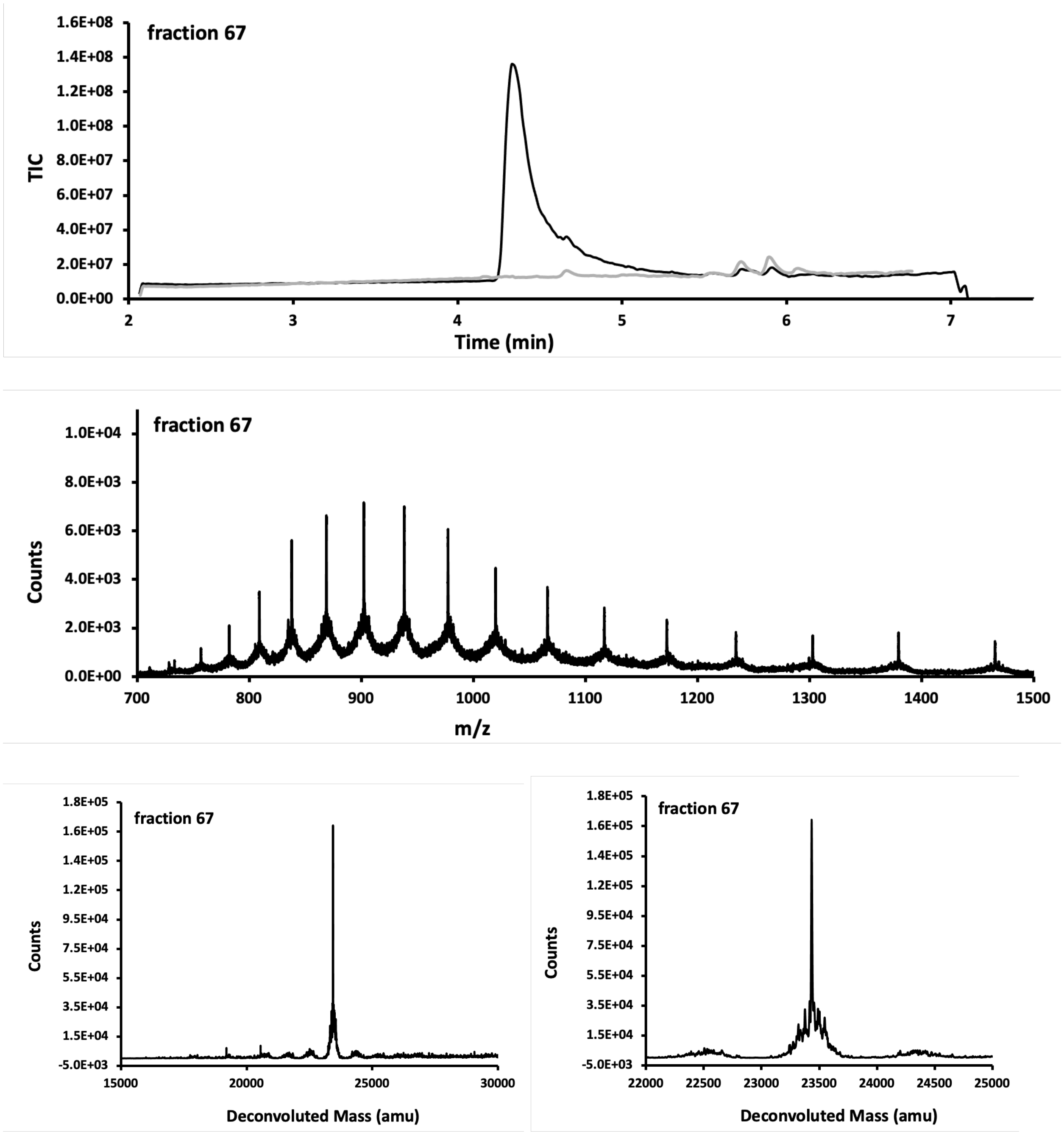


**Figure S16.** LC-MS analysis of fraction 67 of PyS2$^{NTD}$. Top panel is the total ion count versus time. In grey spectrum of a blank run is overlayed. The mass spectrum in the middle panel is obtained from integration over the entire peak and is presented in m/z (in Da) versus time. Bottom panels display deconvoluted mass spectra. This fraction combined to yield the final product. Calc.:23435 Da, Obs. 23423 Da.

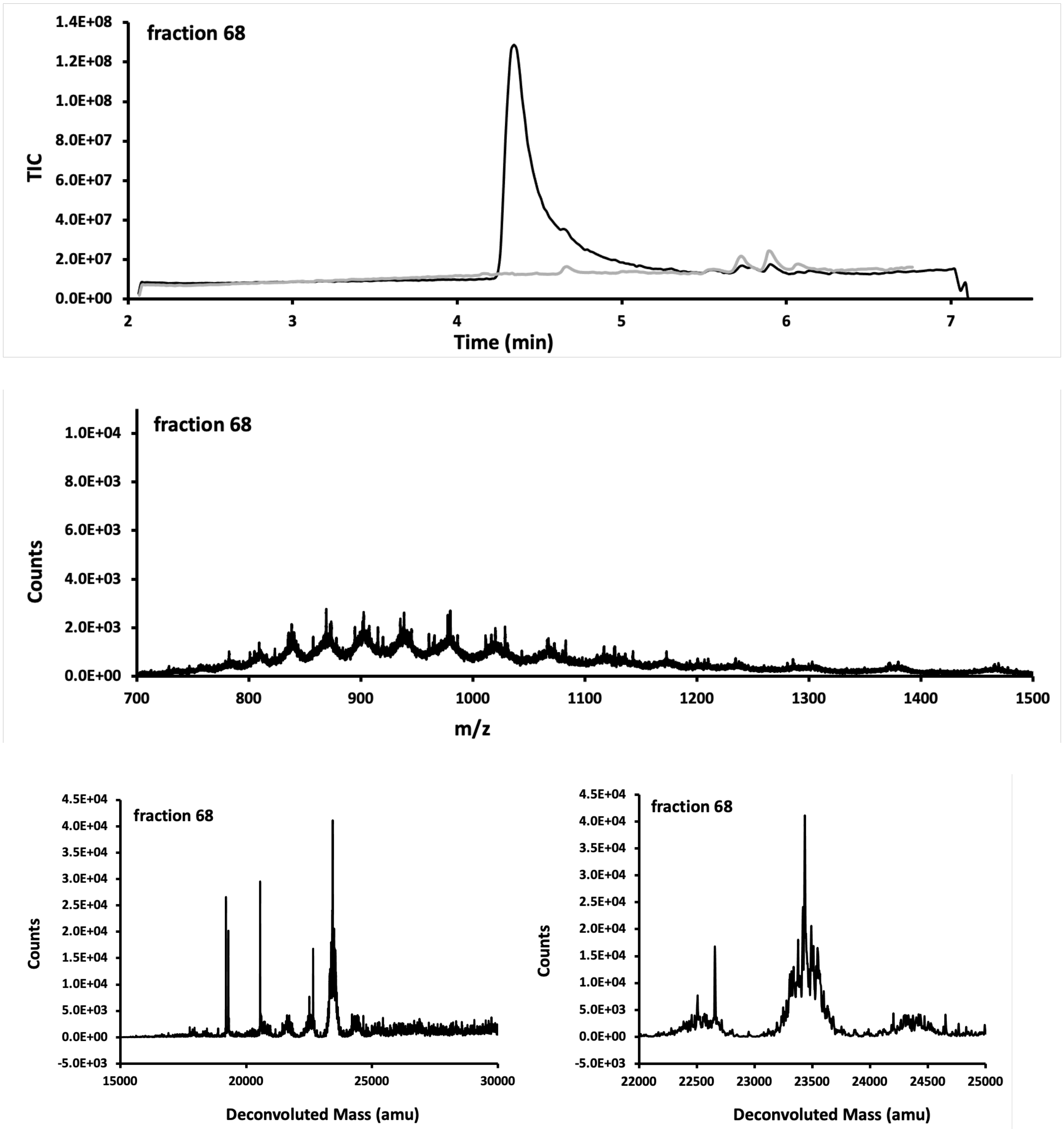


**Figure S17.** LC-MS analysis of fraction 68 of PyS2$^{NTD}$. Top panel is the total ion count versus time. In grey spectrum of a blank run is overlayed. The mass spectrum in the middle panel is obtained from integration over the entire peak and is presented in m/z (in Da) versus time. Bottom panels display deconvoluted mass spectra. This fraction did not exhibit enough purity and was discarded. Calc.:23435 Da, Obs. 23423 Da.

**SI 7.0 Characterization of HPLC-purified PyS2$^{NTD}$**

HPLC-purified PyS2$^{NTD}$ was characterized by HPLC and LC-MS.

For assessment of purity by HPLC, lyophilized HPLC-purified PyS2$^{NTD}$ was dissolved in a freshly prepared solution of 1:1 acetonitrile:water with 0.1% TFA and 500 mM DTT, to yield a solution of PyS2 NTD (at minimum 50 µL) at approximately 1.0 mg/mL. The solution was centrifuged for 2 min at 16873 RCF using a benchtop centrifuge (Eppendorf) and the supernatant was analyzed on an analytical Agilent Zorbax 300SB-C3 column (2.1 mm × 150 mm, 5-µm particle size) on Agilent Technologies 1200 Series instrument, which was computer-controlled through Agilent ChemStation software. A linear gradient of acetonitrile with a 0.08% TFA additive (solvent B) in water with a 0.1% TFA additive (solvent A) was used. Gradients of 5% B per minute ramped up to 65% over 30 min at a flow rate of 0.4 mL/min. Wash and recalibration of the column sequence included ramp up of gradient to 100% B at 31 min, hold at 100% B until 36 min, ramp down to 5% B at 37 min, and hold at 5% B until 43 min. The total method time was 43 min.

For LC-MS method B was used as described in "Analysis method with Liquid-Chromatography Mass Spectrometry (LC-MS)" section.

**SI 7.1. Analytical HPLC of HPLC-purified PyS2$^{NTD}$**

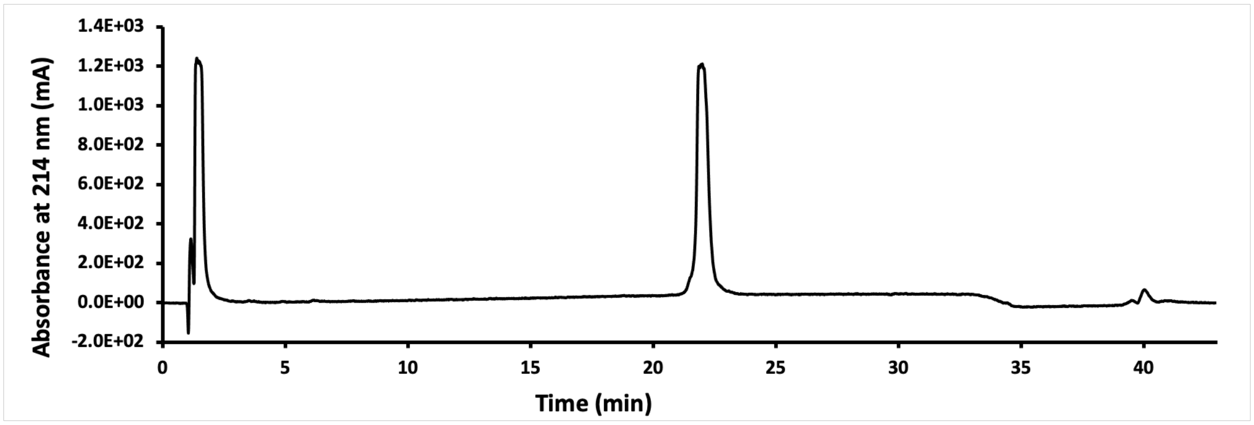


**Figure S18.** HPLC chromatogram of the crude PyS2$^{NTD}$ displaying the absorbance at 214 nm.

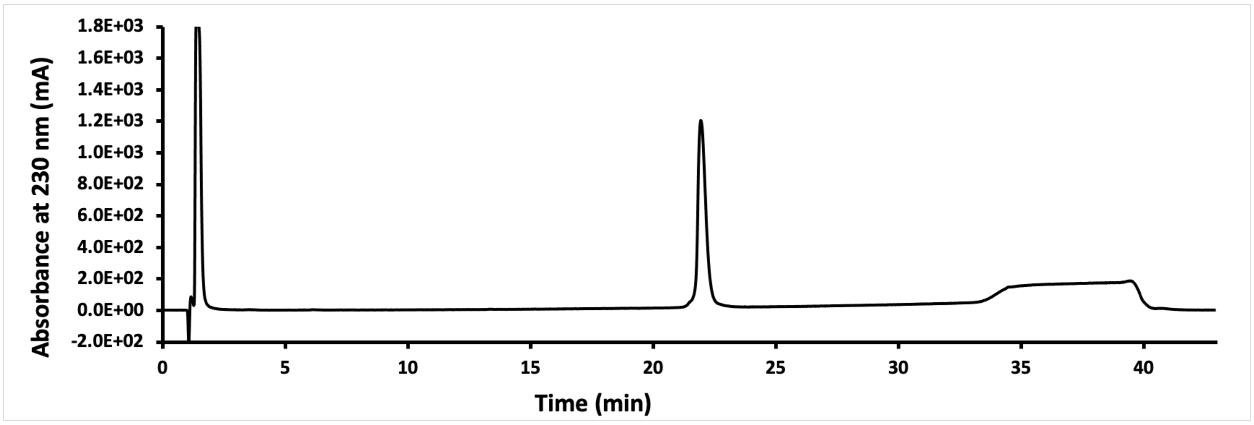


**Figure S19.** HPLC chromatogram of the crude PyS2$^{NTD}$ displaying the absorbance at 230 nm.

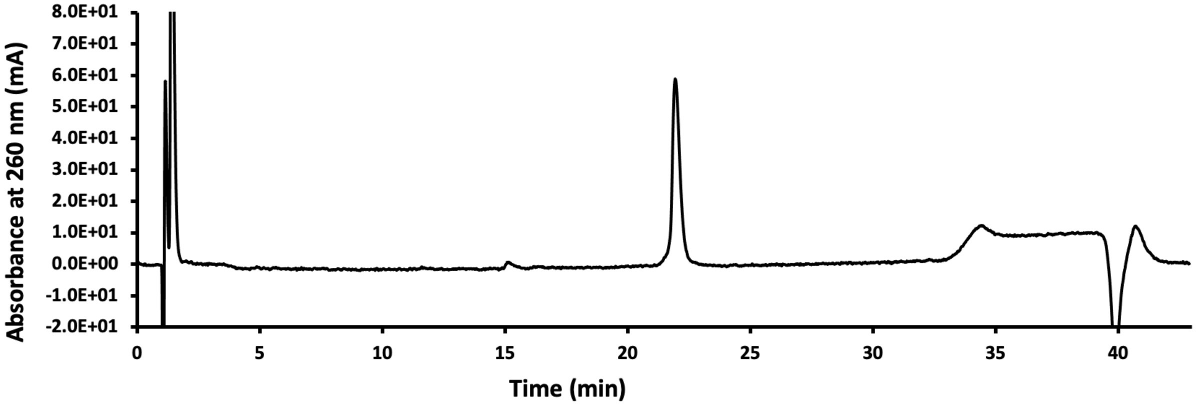


**Figure S20.** HPLC chromatogram of the crude PyS2$^{NTD}$ displaying the absorbance at 260 nm.

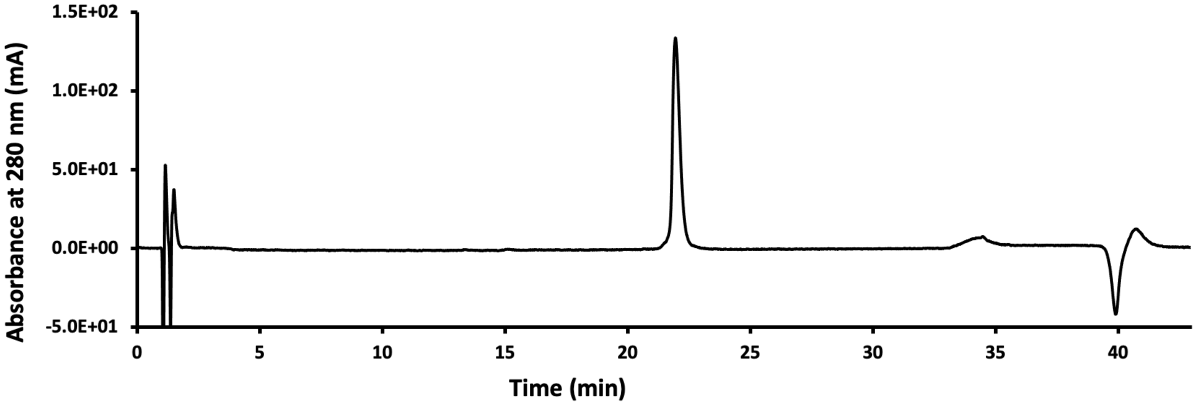


**Figure S21.** HPLC chromatogram of the crude PyS2$^{NTD}$ displaying the absorbance at 280 nm.

### LCMS of HPLC-purified PyS2$^{NTD}$

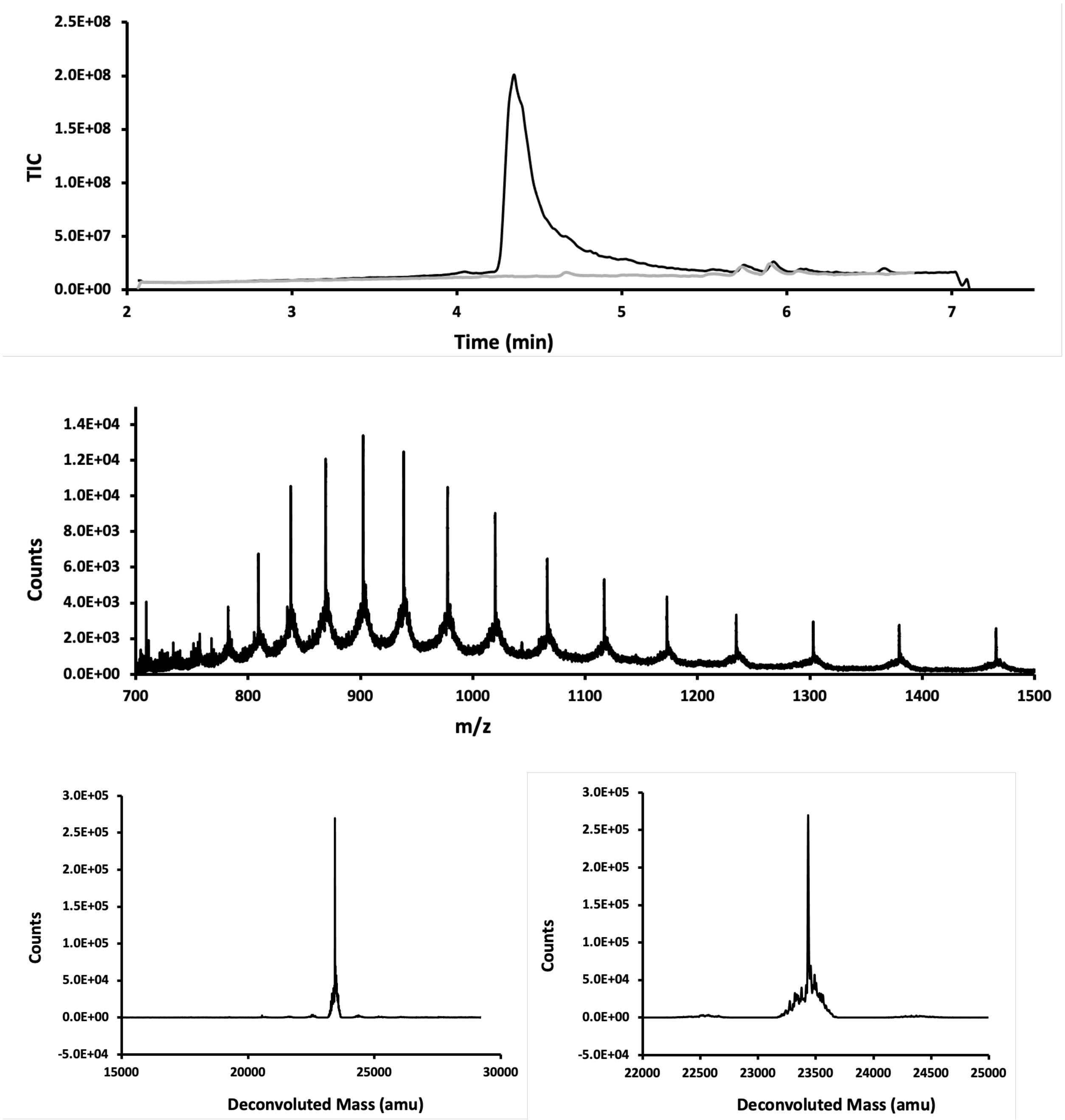


**Figure S22.** LC-MS analysis of HPLC-purified PyS2$^{NTD}$. Top panel is the total ion count versus time. In grey spectrum of a blank run is overlayed. The mass spectrum in the middle panel is obtained from integration over the entire peak and is presented in m/z (in Da) versus time. Bottom panels display deconvoluted mass spectra. Calc.:23435 Da, Obs. 23423 Da.

### SI 8.0. PyS2$^{NTD}$ Folding

A Superdex 10/300 increase 75 column (Cytiva) was equilibrated with 10% v/v glycerol in PBS buffer pH 7.4, using a 1260 Infinity II Bio-inert LC System instrument (Agilent) equipped with a 200 μL loading loop. The column was cleaned prior to folding/purification on reversed flow with 2 column volumes of 0.5 M NaOH or 6 M guanidinium chloride, followed by 1 column volume of water and 3 column volumes of the running buffer (10% v/v glycerol in PBS buffer pH 7.4). A 10 mg/mL solution of PyS2$^{NTD}$ was prepared using HPLC-purified trifluoroacetate salt of PyS2$^{NTD}$ dissolved in 6 M guanidinium chloride in 50 mM $NaH_2PO_4$, pH 7.5, and centrifuged for 2 min at 16873 RCF using a benchtop centrifuge (Eppendorf). Next, 100 μL (36.8 nmol) of the solution was loaded onto the SEC column using the injection loop and eluted using isocratic solution of 10% v/v glycerol in PBS buffer pH 7.4 at 0.5 mL/min flow rate with 0.25 min interval fraction collection. The elution was monitored for changes in absorbance at 214 nm, 230 nm and 280 nm, as well as changes in fluorescence signal with excitation at 280 nm and emittance at 325 nm.

Similar to RP-HPLC purification of PyS2$^{NTD}$, fractions from SEC purification of PyS2$^{NTD}$ were analyzed by LC-MS and high-quality fractions were combined to afford the folded PyS2$^{NTD}$. Overall, 1.5 mL of PyS2$^{NTD}$ was isolated. Concentration of the solution was determined by absorbance at 280 ($A_{280}$) as described in the "determination of folded PyS2$^{NTD}$ concentrations" section and was calculated to be 20.7 μM, corresponding to 0.72 mg of protein (31 nmol, 84% yield).

**Determination of folded PyS2$^{NTD}$ concentrations**

Molar extinction coefficients of folded PyS2$^{NTD}$ was estimated using ExPASy SIB Bioinformatics Resource Portal, which was determined to be $\epsilon_{280}$ = 16,390 $M^{-1}cm^{-1}$

This molar extinction coefficient was used in combination with absorbance at 280 nm ($A_{280}$) in aqueous buffer to determine protein concentration according to Beer's law. The blank for measurement of $A_{280}$ of each sample of PyS2$^{NTD}$ was adjusted accordingly. For example, 10% v/v glycerol in PBS buffer pH 7.4 was used for folded PyS2$^{NTD}$, while 10 mM potassium phosphate pH 7.5 was used for the CD sample. Note that concentration of PyS2-488 was determined by fluorescence, not absorbance.

### SI 8.1. SEC chromatogram of folding of PyS2$^{NTD}$

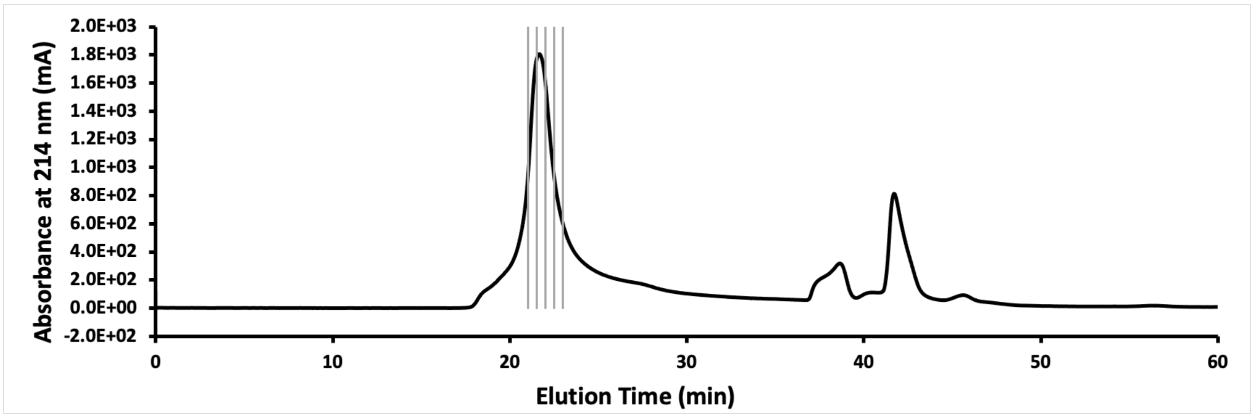


**Figure S23.** Chromatogram from folding and purification of PyS2$^{NTD}$ with SEC displaying the absorbance at 214 nm. Product elutes around the 22-minute mark. Fractions containing the product, as determined by LC-MS, shown in gray, were combined to afford folded PyS2$^{NTD}$ at 84% yield.

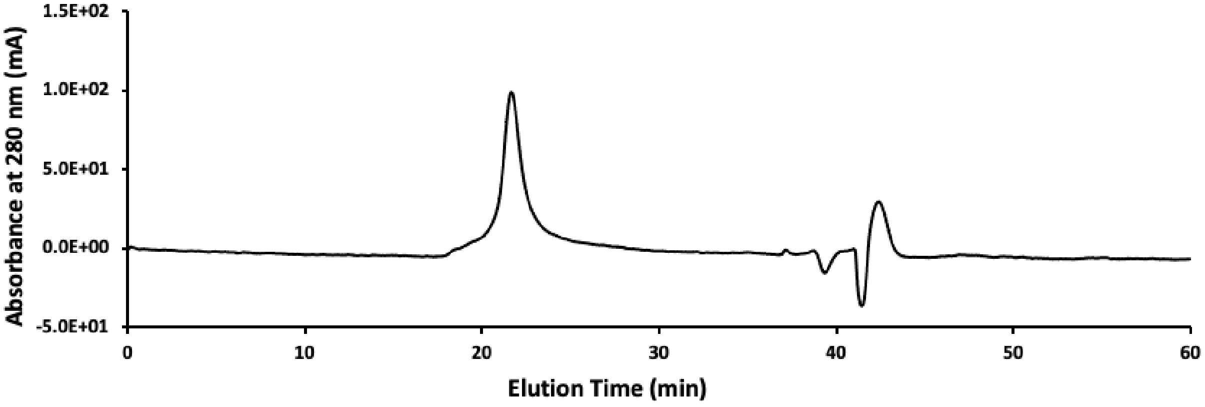


**Figure S24.** Chromatogram from SEC of PyS2$^{NTD}$ displaying the absorbance at 280 nm.

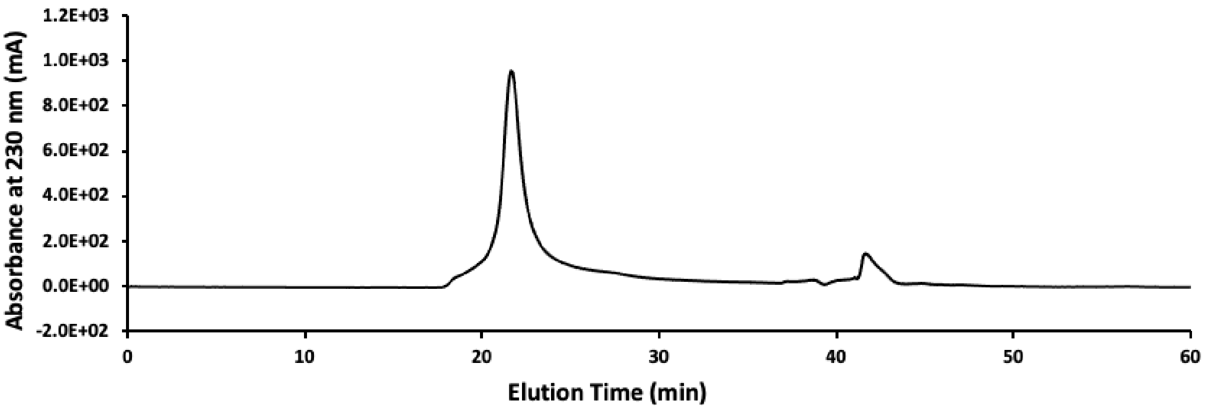


**Figure S25.** Chromatogram from SEC of PyS2$^{NTD}$ displaying the absorbance at 230 nm.

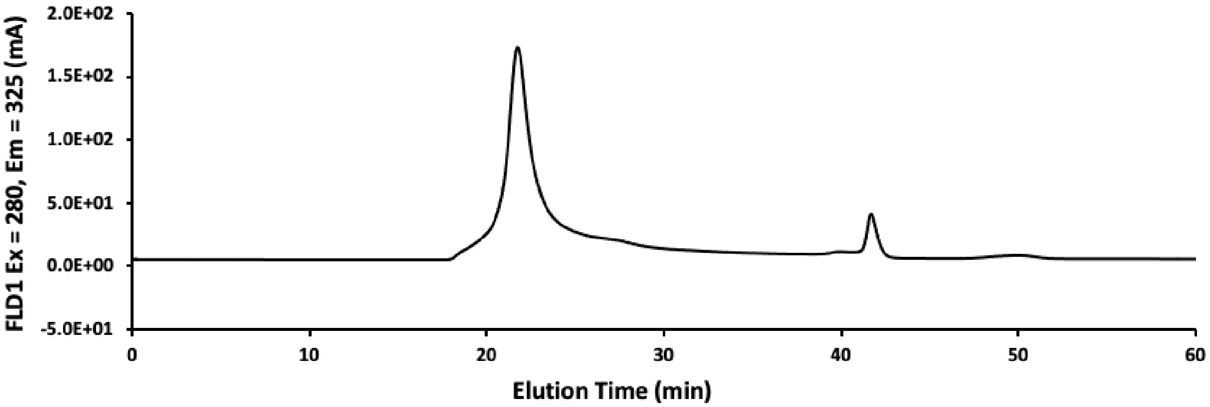


**Figure S26.** Chromatogram from SEC of PyS2$^{NTD}$ displaying the fluorescence signal with excitation at 280 nm and emission at 325 nm.

### SI 9.0. Characterization of folded PyS2$^{NTD}$

Folded PyS2$^{NTD}$ was characterized by HPLC and LC-MS.

For assessment of purity by HPLC, the solution of the PyS2$^{NTD}$ was diluted to ~0.5 mg/mL using a solution of 1:1 acetonitrile:water with 0.1% TFA and 500 mM DTT. The solution was then analyzed on an analytical Agilent Zorbax 300SB-C3 column (2.1 mm × 150 mm, 5-µm particle size) on Agilent Technologies 1200 Series instrument, which was computer-controlled through Agilent ChemStation software. A linear gradient of acetonitrile with a 0.08% TFA additive (solvent B) in water with a 0.1% TFA additive (solvent A) was used. Gradients of 5% B per minute ramped up to 65% over 30 min at a flow rate of 0.4 mL/min. Wash and recalibration of the column sequence included ramp up of gradient to 100% B at 31 min, hold at 100% B until 36 min, ramp down to 5% B at 37 min, and hold at 5% B until 43 min. The total method time was 43 min.

For LC-MS method B was used as described in "Analysis method with Liquid-Chromatography Mass Spectrometry (LC-MS)" section.

**SI 9.1. Analytical HPLC of folded PyS2$^{NTD}$**

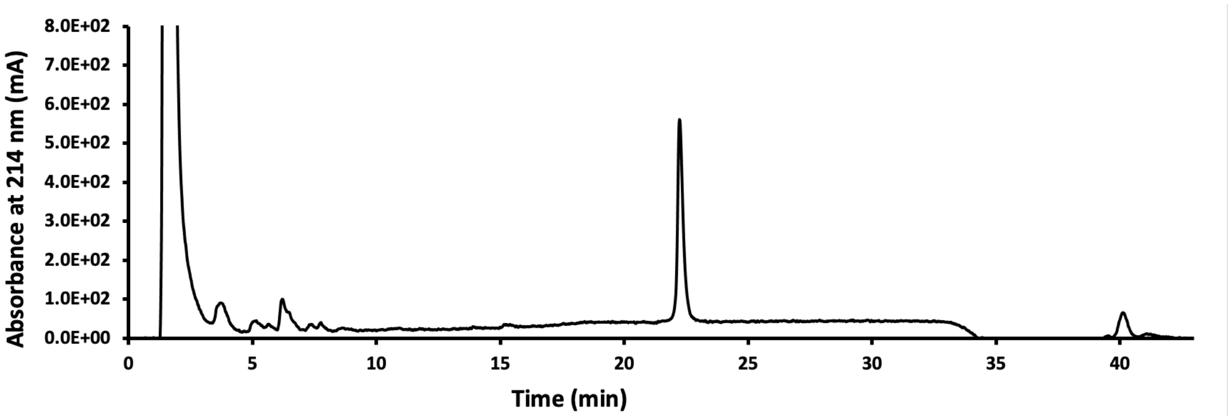


**Figure S27.** HPLC chromatogram of the folded PyS2$^{NTD}$ displaying the absorbance at 214 nm.

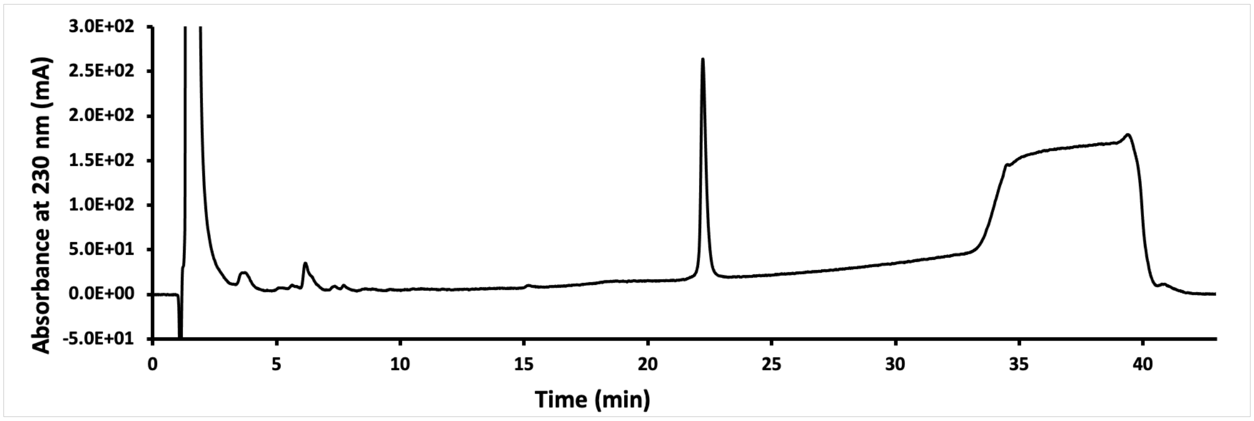


**Figure S28.** HPLC chromatogram of the folded PyS2$^{NTD}$ displaying the absorbance at 230 nm.

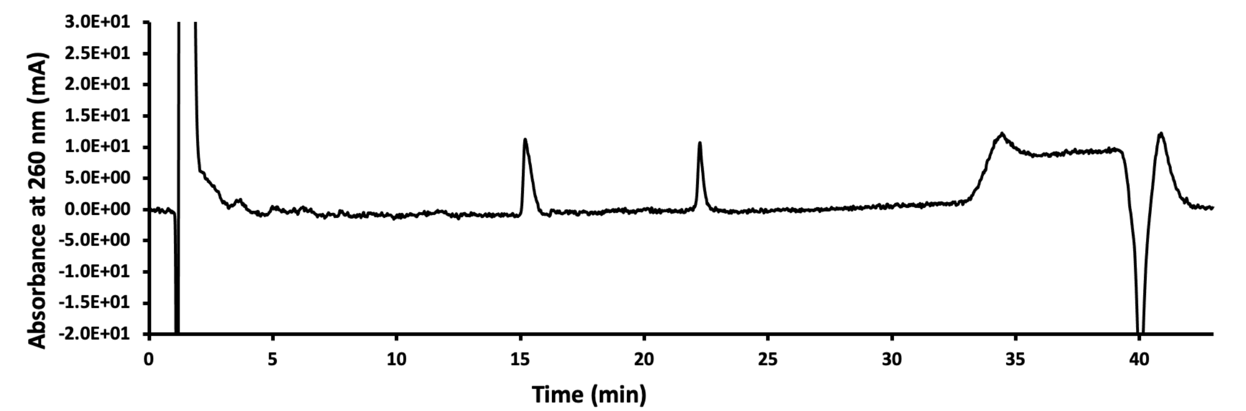


**Figure S29.** HPLC chromatogram of the folded PyS2$^{NTD}$ displaying the absorbance at 260 nm.

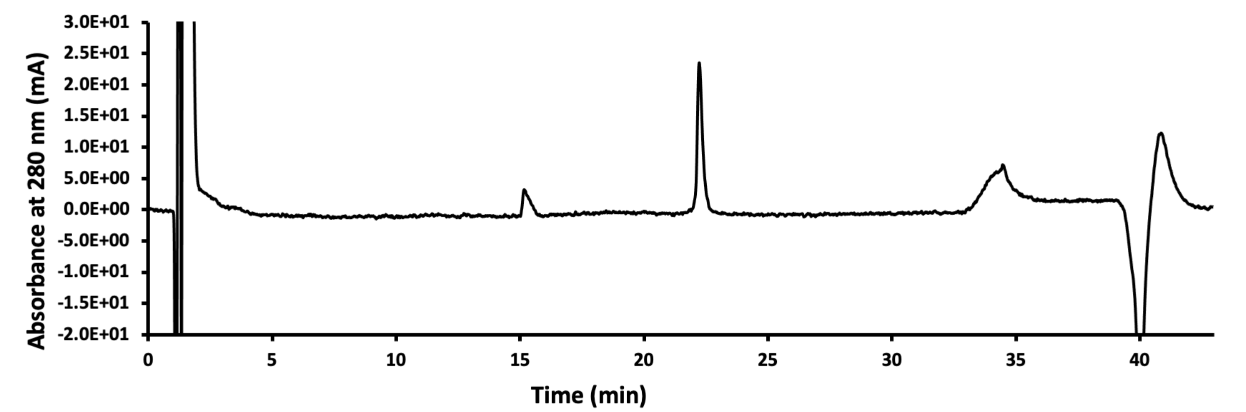


**Figure S30.** HPLC chromatogram of the folded PyS2$^{NTD}$ displaying the absorbance at 280 nm.

### SI 9.2. LC-MS of folded PyS2$^{NTD}$

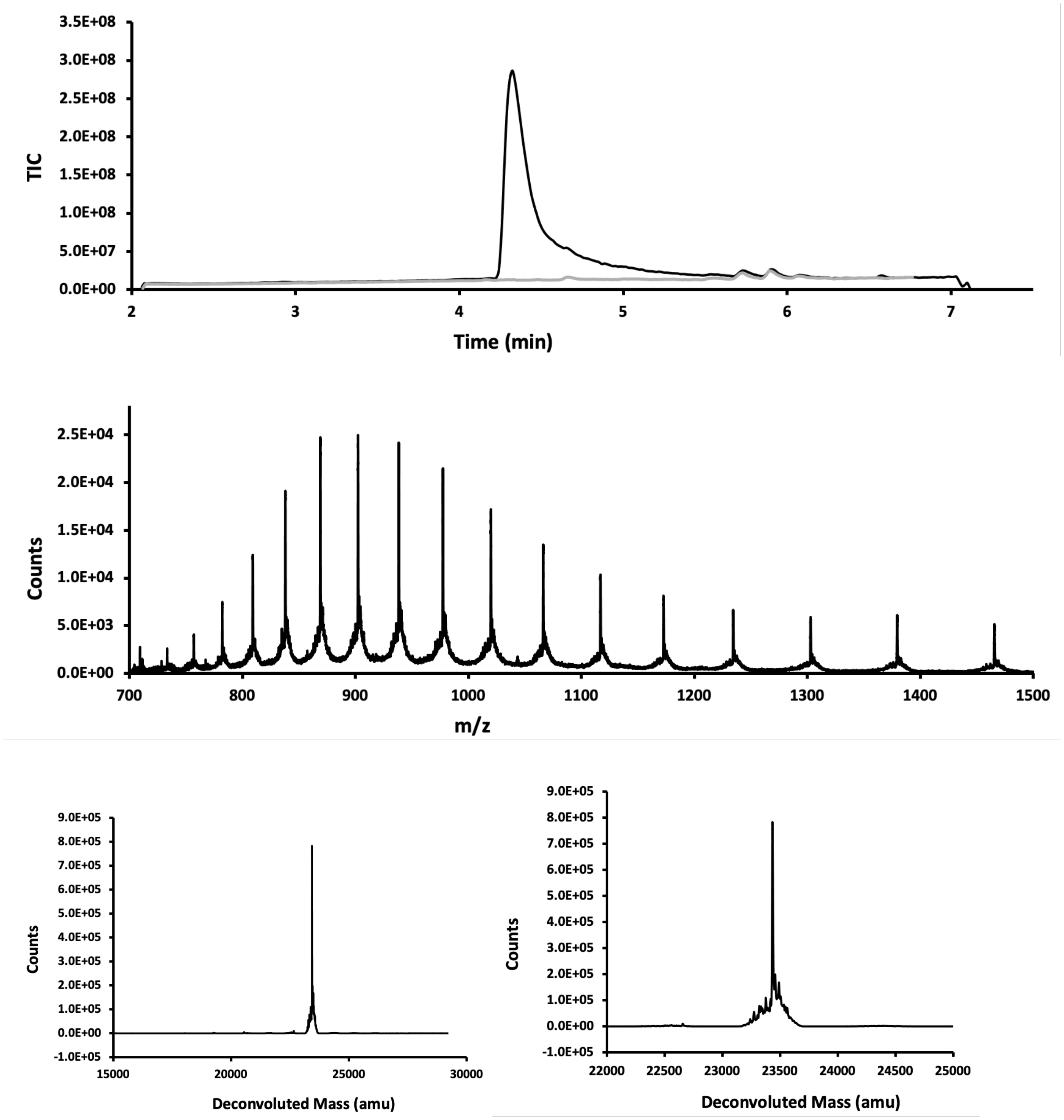


**Figure S31.** LC-MS analysis of folded PyS2$^{NTD}$. Top panel is the total ion count versus time. In grey spectrum of a blank run is overlayed. The mass spectrum in the middle panel is obtained from integration over the entire peak and is presented in m/z (in Da) versus time. Bottom panels display deconvoluted mass spectra. Calc.:23435 Da, Obs. 23423 Da.

**SI 10.0. Circular dichroism measurements**

PyS2$^{NTD}$ was buffer exchanged into 10 mM potassium phosphate pH 7.5 using 3K molecular weight cut off spin filters. The concentration of the sample was measured based on the absorbance of the sample at 280 nm.

Spectra were collected from 196-260 nm (1 nm step size, 0.33 second settling time, 1 nm monochromator bandwidth) on an AVIV Biomedical Circular Dichroism Spectrometer Model 420 (Lakewood, NJ) using a 0.1 cm cuvette filled with 350 uL of PyS2$^{NTD}$ at 0.1 mg/mL in 10 mM phosphate buffer, pH 7.5 at 25 °C. The CD signal (mdeg) was converted to mean residue ellipticity (MRE) using the length, concentration, and molecular weight of the protein.[2] A thermal melt was completed by incrementing the temperature of the sample from 25 °C to 70 °C with a minimum of 2 minutes equilibration once the setpoint temperature was achieved using an automated script to gather spectra. The thermal melt was calculated by fitting the data to the model of a two-state transition of a monomer from a folded to unfolded state.[3] Data was plotted in GraphPad Prism, including smoothing (0$^{th}$ order, 2 neighbors to average) and nonlinear fitting of the thermal melt model.

### SI 11.0. Fluorophore conjugation of PyS2$^{NTD}$

**Reaction conditions.** In a 1 mL conical Eppendorf tube 1.17 mg HPLC-purified PyS2$^{NTD}$ (43 nmol, 1 equiv) was added and dissolved in 94.8 μL 6 M guanidinium chloride. Next, 5.2 μL of 10 mM solution of AZDye 488 Maleimide (Fluoroprobes, cat#1014-1) in DMSO was added to the mixture (52 nmol, 1.2 equiv). The tube was covered with aluminum foil to block light and left at room temperature for an hour. Following confirmation that the PyS2$^{NTD}$ is fully used up in the reaction by LC-MS (method C, product displays +699 Da in mass, Figure S30), the mixture was loaded on a Superdex 10/300 increase 75 column using a 200 mL loading loop and eluted with isocratic solution of 10% v/v glycerol in PBS pH 7.4 as the mobile phase. The column was cleaned prior to purification on reversed flow with 2 column volumes of 0.5 M NaOH or 6 M guanidinium chloride, followed by 1 column volume of water and 3 column volumes of the running buffer (10% v/v glycerol in PBS buffer pH 7.4). Fractions were collected and analyzed by LC-MS. The Fractions containing the product and showing high-purity charge state series were combined to afford 0.234 mg PyS2-488 (10 nmol) in 24% yield.

#### Determination of PyS2-488 concentrations

Concentration of PyS2-488 was determined by measuring the fluorescence intensity of the PyS2-488 sample. A standard curve of the AZDye 488 Maleimide was created by serial dilution in 10% v/v glycerol in PBS buffer pH 7.4. Five samples of the dye with concentrations of 50 nM, 20 nM, 10 nM, 5 nM, 2 nM were used to generate the standard curve. Samples were made in triplicates and their fluorescence intensity were measured in triplicates using a Tecan plate reader M1000 with excitation at 483 nm and emission at 528 nm. Within the same run, three dilutions of the purified PyS2-488 solution were also measured for their fluorescence intensity in triplicates.

Specifically, 300-fold diluted-, 500-fold diluted and 1000 -old diluted-PyS2-488 were used. Using the standard curve, in this case the concentration of the PyS2-488 was determined to be 10 μM.

| concentration (nM) | measured fluorescence intensity |
|---|---|
| 50 | 47,562 |
| 20 | 19,330 |
| 10 | 10,083 |
| 5 | 5,079 |
| 2 | 2,022 |

| Dilution factor of PyS2-488 | measured fluorescence intensity |
|---|---|
| 300 | 32,696 |
| 500 | 19,369 |
| 1000 | 9,617 |

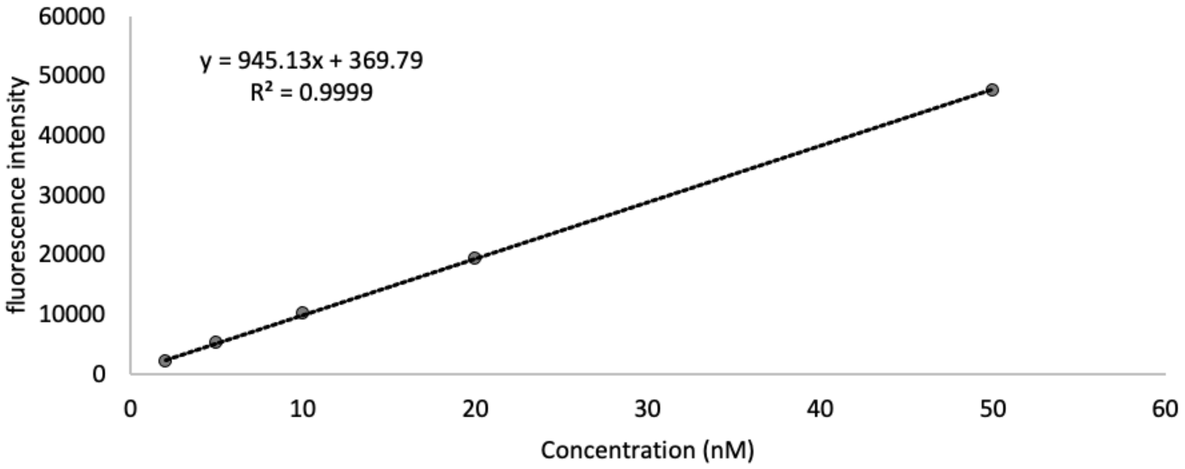


**Figure S32.** Standard curve generated with dilutions of AZDye 488 Maleimide in 10% v/v glycerol in PBS buffer pH 7.4 for measuring the concentration of the PyS2-488 samples.

### SI 11.1. Monitoring reaction by LC-MS

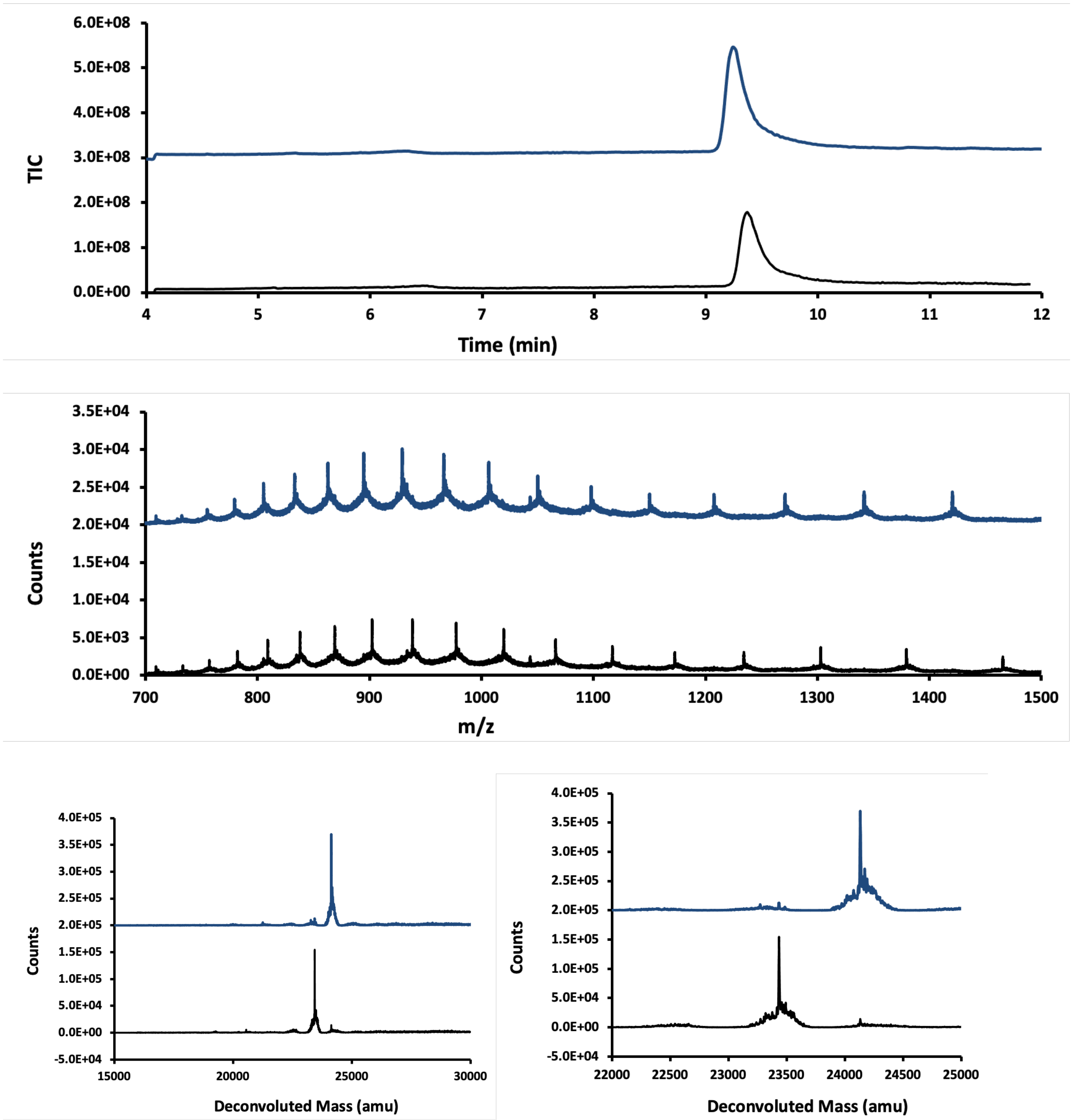


**Figure S33.** LC-MS analysis of PyS2$^{NTD}$ (in black/bottom) and reaction mixture of PyS2$^{NTD}$ with AZDye 488 Maleimide (in blue/top). Top panel is the total ion count versus time. The mass spectrum in the middle panel is obtained from integration over the entire peak and is presented in m/z (in Da) versus time. Bottom panels display deconvoluted mass spectra. Bottom mass: Calc.:23435 Da, Obs. 23423 Da. Top mass: Calc.:24134 Da (+699 Da), Obs. 24234 Da.

**SI 11.2. SEC chromatogram of purification/folding of PyS2-488**

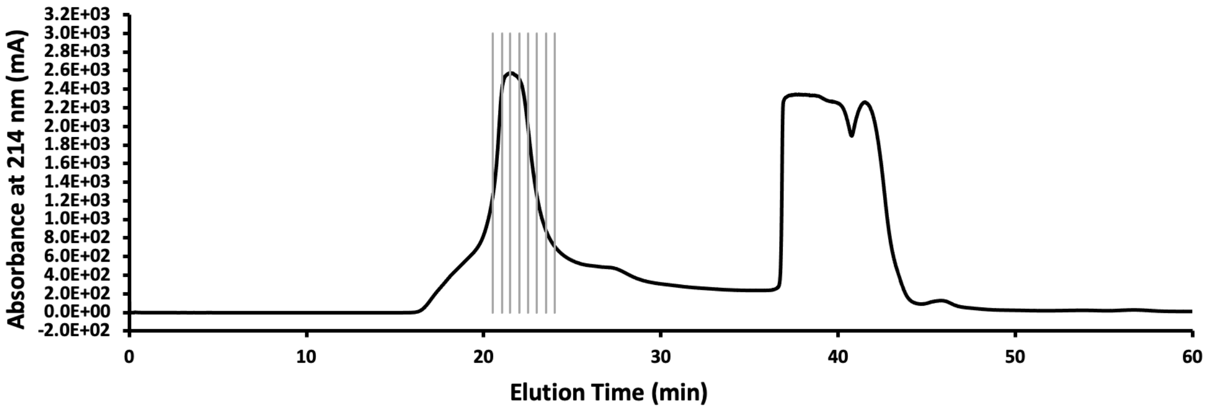


**Figure S34.** Chromatogram from SEC of reaction mixture of PyS2$^{NTD}$ with AZDye 488 Maleimide displaying the absorbance at 214 nm. Fractions containing the product, as determined by LC-MS, shown in gray, were combined to afford folded PyS2$^{NTD}$ at 24% yield.

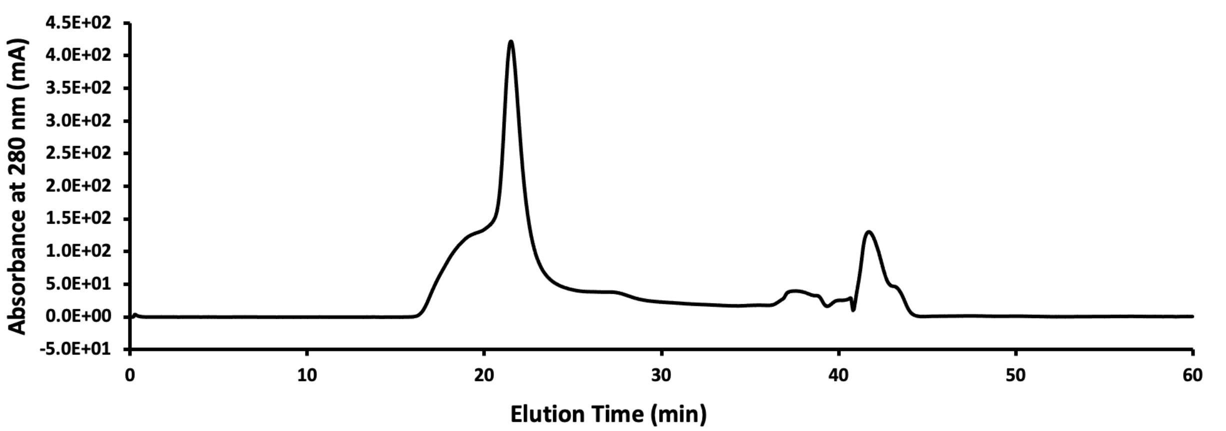


**Figure S35.** Chromatogram from SEC of reaction mixture of PyS2$^{NTD}$ with AZDye 488 Maleimide displaying the absorbance at 280 nm.

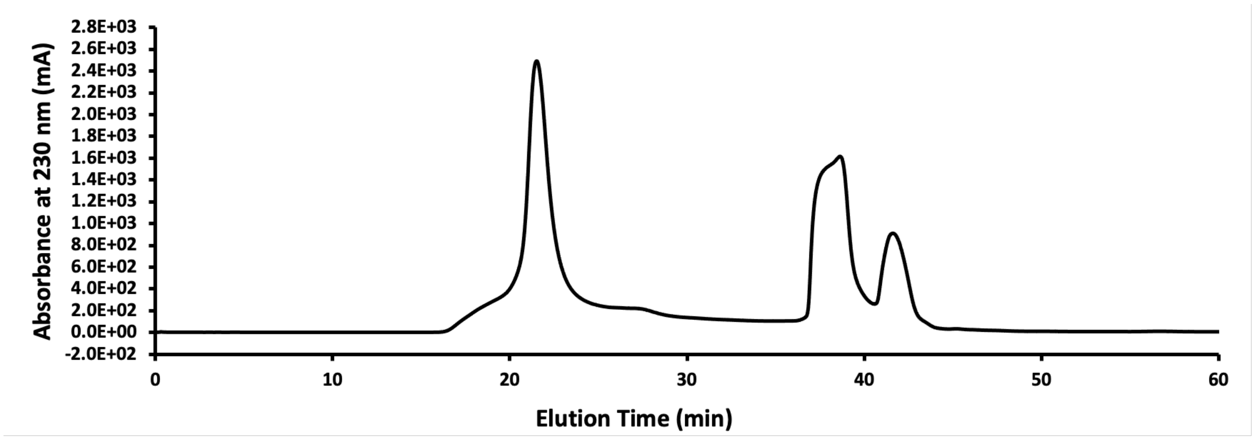


**Figure S36.** Chromatogram from SEC of reaction mixture of PyS2$^{NTD}$ with AZDye 488 Maleimide displaying the absorbance at 230 nm.

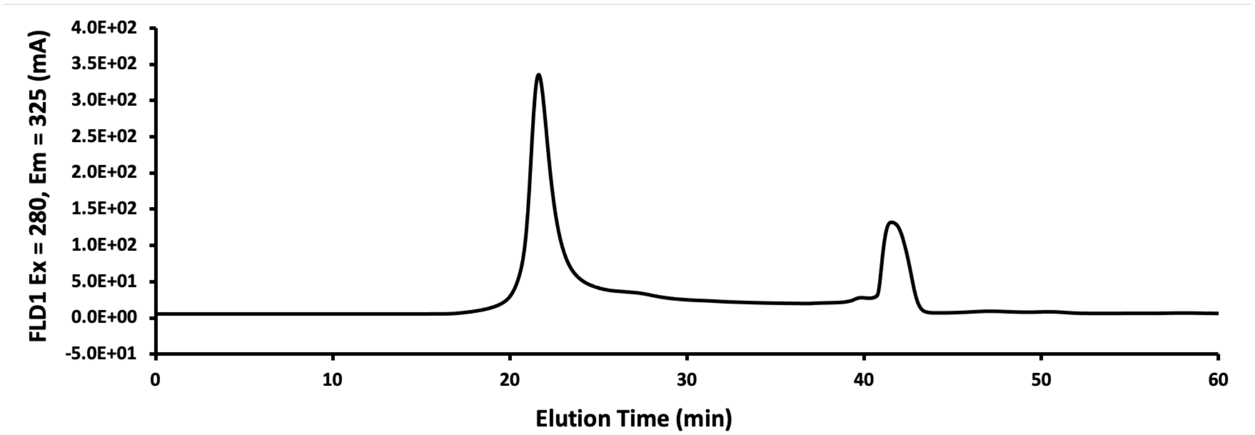


**Figure S37.** Chromatogram from SEC of reaction mixture of PyS2$^{NTD}$ with AZDye 488 Maleimide displaying the fluorescence signal with excitation at 280 nm and emission at 325 nm.

**SI 12.0. Characterization of PyS2-488**

Folded PyS2-488 was characterized by HPLC and LC-MS.

For assessment of purity by HPLC, the solution of the PyS2-488 was diluted to ~0.5 mg/mL using a solution of 1:1 acetonitrile:water with 0.1% TFA. The solution was then analyzed on an analytical Agilent Zorbax 300SB-C3 column (2.1 mm × 150 mm, 5-µm particle size) on Agilent Technologies 1200 Series instrument, which was computer-controlled through Agilent ChemStation software. A linear gradient of acetonitrile with a 0.08% TFA additive (solvent B) in water with a 0.1% TFA additive (solvent A) was used. Gradients of 5% B per minute ramped up to 65% over 30 min at a flow rate of 0.4 mL/min. Wash and recalibration of the column sequence included ramp up of gradient to 100% B at 31 min, hold at 100% B until 36 min, ramp down to 5% B at 37 min, and hold at 5% B until 43 min. The total method time was 43 min.

For LC-MS method C was used as described in "Analysis method with Liquid-Chromatography Mass Spectrometry (LC-MS)" section.

### SI 12.1. Analytical HPLC analysis of PyS2-488

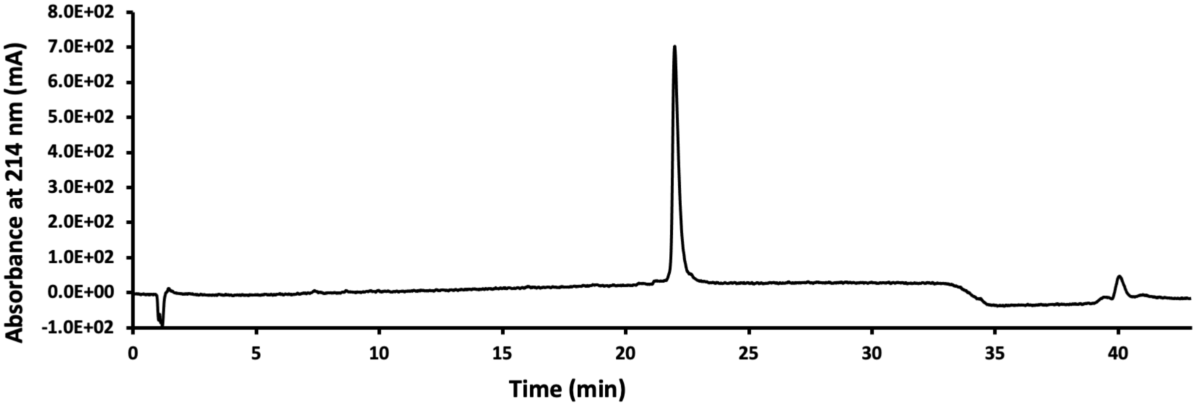


**Figure S38.** HPLC chromatogram of the PyS2-488 displaying the absorbance at 214 nm.

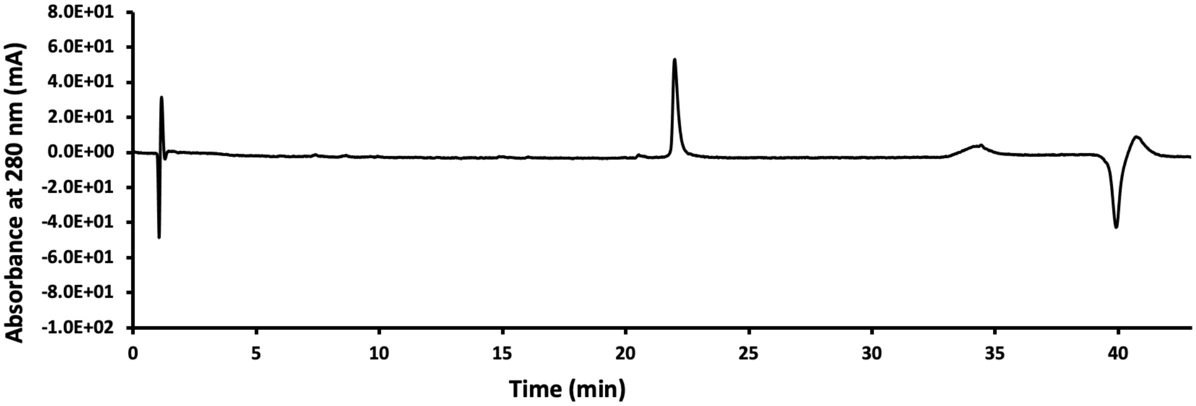


**Figure S39.** HPLC chromatogram of the PyS2-488 displaying the absorbance at 280 nm.

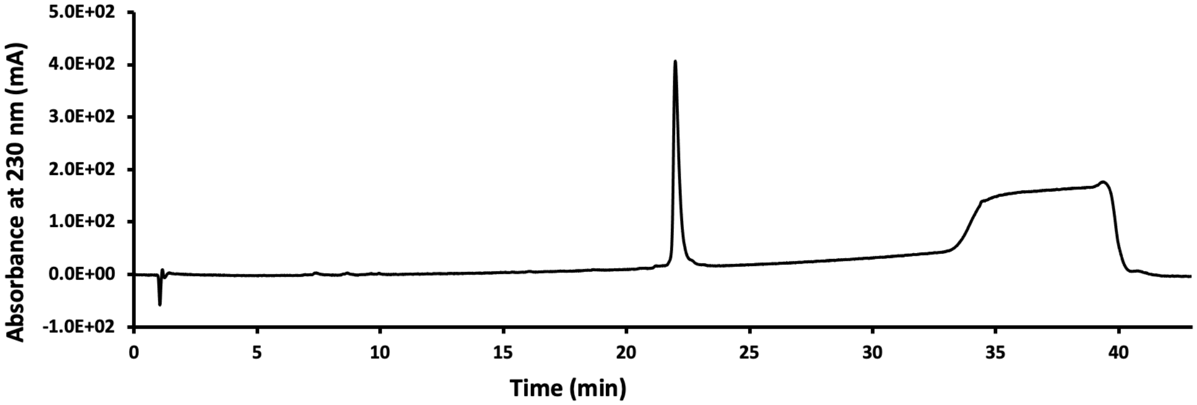


**Figure S40.** HPLC chromatogram of the PyS2-488 displaying the absorbance at 230 nm.

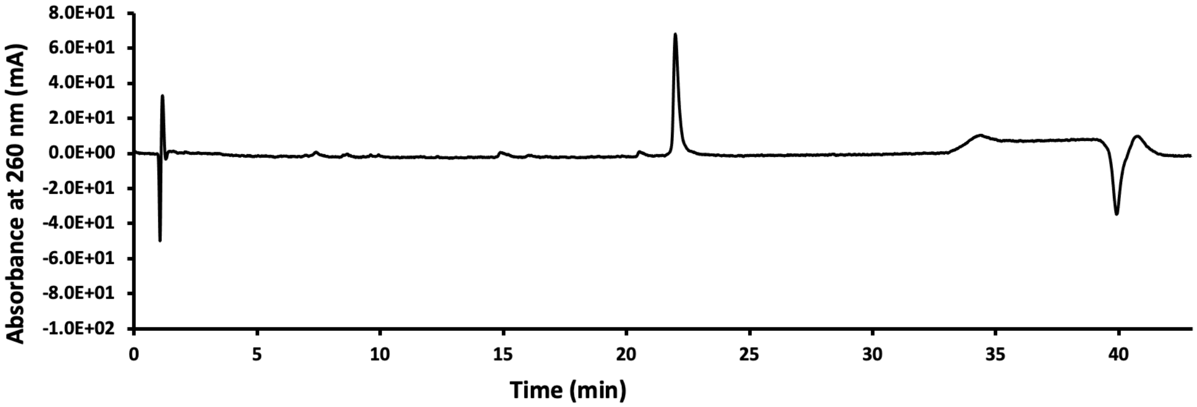


**Figure S41.** HPLC chromatogram of the PyS2-488 displaying the absorbance at 260 nm.

### SI 12.2. LC-MS of PyS2-488

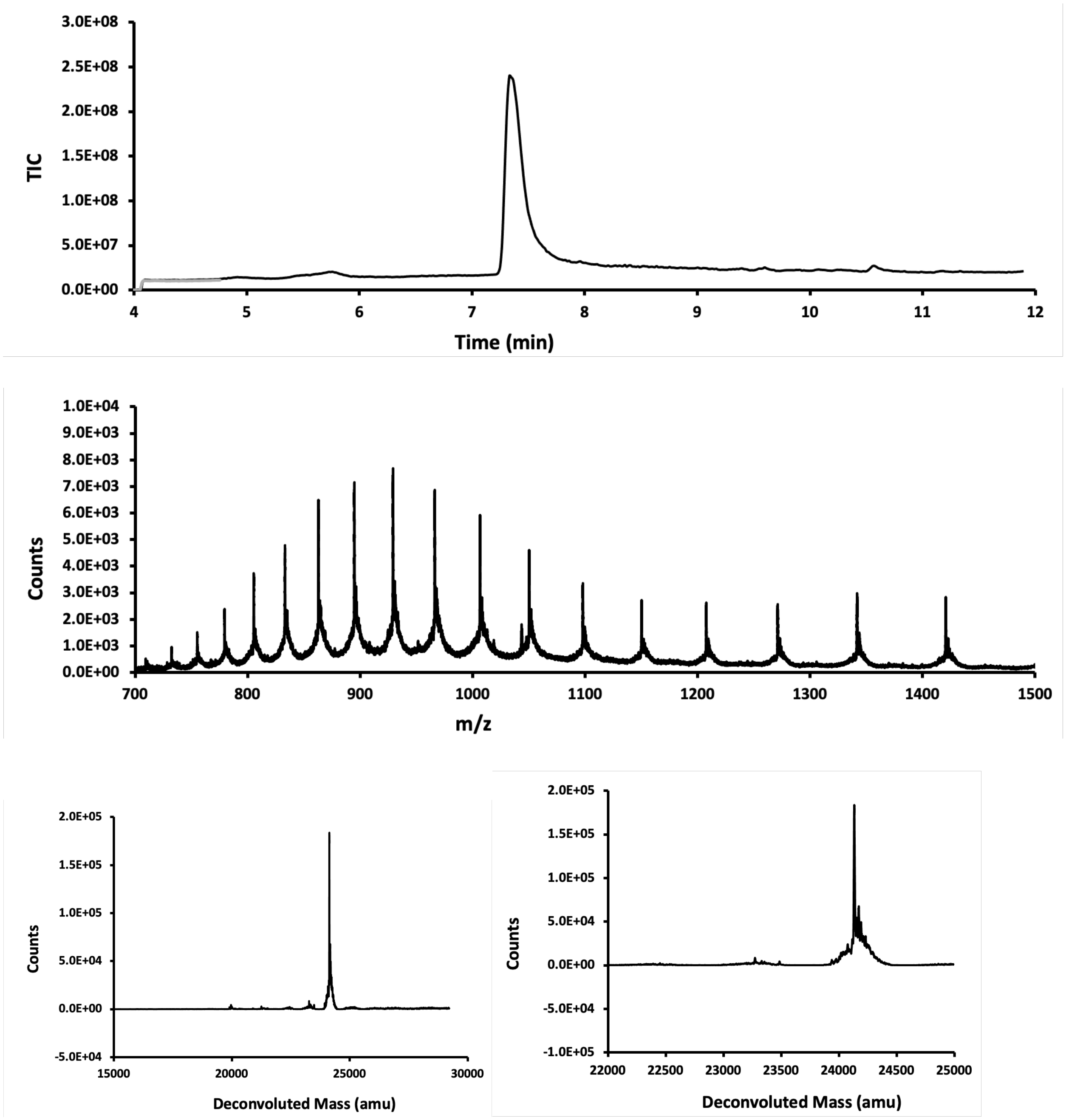


**Figure S42.** LC-MS analysis of PyS2-488. Top panel is the total ion count versus time. In grey spectrum of a blank run is overlayed. The mass spectrum in the middle panel is obtained from integration over the entire peak and is presented in m/z (in Da) versus time. Bottom panels display deconvoluted mass spectra. Calc.: 24134 Da, Obs. 24134 Da.

### SI 13.0. Biological Characterization of PyS2$^{NTD}$

#### SI 13.1. Bacterial Strains and Construction

PAO1 Δ*fpvA* was constructed as previously described.[4] Briefly, the flanking genes were amplified, SOEn together, and inserted into a pEXG2 vector using gateway sites. Merodiploid colonies were allowed to recombine and second recombination events were selected for using sucrose and selection antibiotic. Clean deletion of *fpvA* was confirmed by PCR amplification and sequencing.

#### SI 13.2. Staining of *Pseudomonas aeruginosa* with PyS2$^{NTD}$-fluorophore conjugate

*E. coli* (ATCC 25922), PAO1, and *ΔfpvA* were grown overnight in Luria broth (LB), M9 glucose, or M9 glucose supplemented with 17 µM ferric (+3) iron chloride at 37 °C to stationary phase. The cultures were spun for 10 minutes at 3.2 k rcf, resuspended and washed with 0.5% BSA in M9 glucose media (M9G) pH 7.4 with 0.01% Tween80 (blocking buffer) three times. Bacteria were stained with 1 µM PyS2$^{NTD}$-fluorophore conjugate and 5 µM cell-permeant SYTO 40 Blue at an OD600 of 3.6 in 100 µL for 30 minutes at room temperature and then washed three times again with blocking buffer. For samples where internalization was inhibited with CCCP, 100 µM CCCP was used 10 minutes before staining and maintained throughout all washes. Similarly, 5 µM SYTO 40 Blue was maintained in all washes. After the last wash, cells were resuspended in 0.5% BSA in M9G with 0.05% Tween80 with 5 µM SYTO 40 Blue.

**SI 13.3. Flow cytometry**

Stained and unstained bacteria-only controls were diluted ten-fold in 0.5% BSA in M9G with 0.05% Tween80 with 5 μM SYTO 40 Blue and placed into flow tubes (Corning Cat. 352235), if present, 100 μM of CCCP was maintained in samples. Samples were then analyzed using an Attune NxT Flow Cytometer with 405 nm, 488 nm, 561 nm, and 640 nm lasers. 10,000 cells were counted at the low flow rate. Flow cytometry analysis was performed in biological triplicate to gather experimental error bars. The unstained controls were analyzed first to set gates. Data were analyzed using the FlowJo software package (FlowJo LLC). Mean fluorescence intensity was calculated using a geometric mean.

**SI 13.4. Confocal microscopy and fluorescence recovery after photobleaching (FRAP)**

Stained and unstained bacteria-only controls in 0.5% BSA in M9G with 0.05% Tween80 with 5 μM SYTO 40 Blue, and if present, 100 μM CCCP were suspended. Bacteria suspensions were dispensed (3-5 μL) on to pre-prepared pads of 1% agarose in M9G (with CCCP if needed) assembled using a 125 μL (1.7 x 2.8 cm) Gene Frame on a plain 1.0 mm glass microscope slide (2.5 x 7.5 cm) and then sealed with a coverslip (thickness 1 1/2, 2.4 x 40 cm). Samples were then visualized at the W. M. Keck Microscopy Facility at the Whitehead Institute using an Andor Revolution Spinning Disk Confocal using a 100× oil immersion objective with 1.4 N.A. Differential Interference Contrast (brightfield) images were collected. Fluorescent confocal images were taken by exciting at 405 nm and 488 nm to observe the SYTO 40 Blue at 447/60 nm and AzDye488 (PyS2$^{NTD}$-fluorophore conjugate) at 525/40 nm, respectively on a Andor iXon 897E back-illuminated EM-CCD camera. FRAP was completed using the Andor FRAPPA photomanipulation system (A portion of each cell was photobleached with a dwell time of 250 μs, twice) to FRAP multiple bacteria (10-15) in a single field of view. Images were acquired before

FRAP and recovery was observed afterward every 10 seconds for 2 minutes. Images were processed using ImageJ. Recovery was calculated by normalizing using the following equation as previously reported using the equation below.[5] The values were initialized at 1 and immediately after photobleaching, the value was 0.

$$F(t)_{\text{norm}} = \frac{F(t)_{\text{ROI}} - F_{\text{bkgd}}}{F(t)_{\text{cell}} - F_{\text{bkgd}}} \times \frac{F(i)_{\text{cell}} - F_{\text{bkgd}}}{F(i)_{\text{ROI}} - F_{\text{bkgd}}}$$

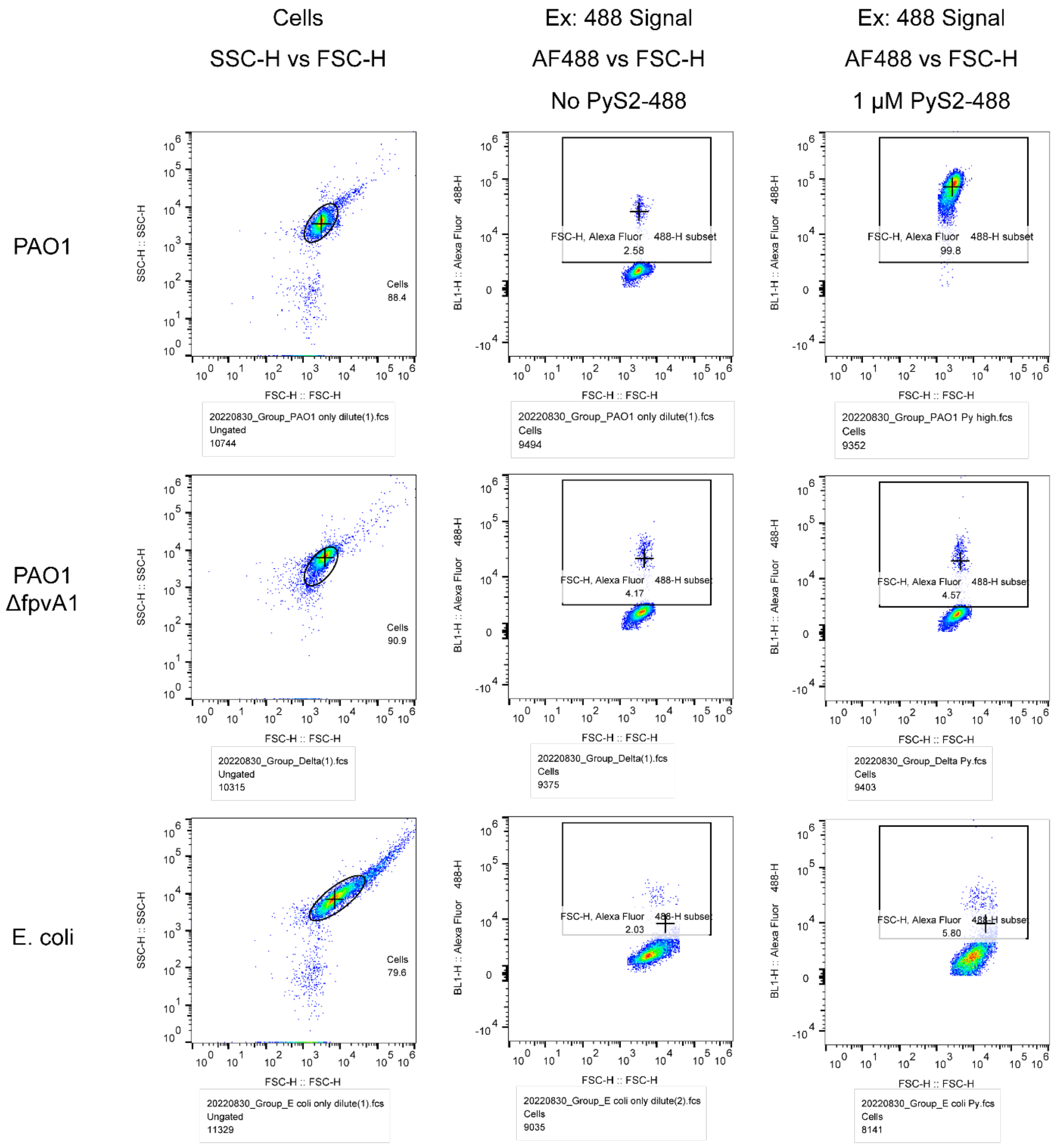


**Figure S43.** Raw representative flow cytometry plots from Main Text Figure 4 indicating species-specific binding of PyS2-488 in comparison to PAO1 Δ*fpvA* and *E. coli* (ATCC 25922) when grown in iron-poor minimal M9 Glucose (M9G) media. 10,000 events were recorded with the final gate number of events listed below each graph. Each graph is a representative of triplicate biologic experiments.

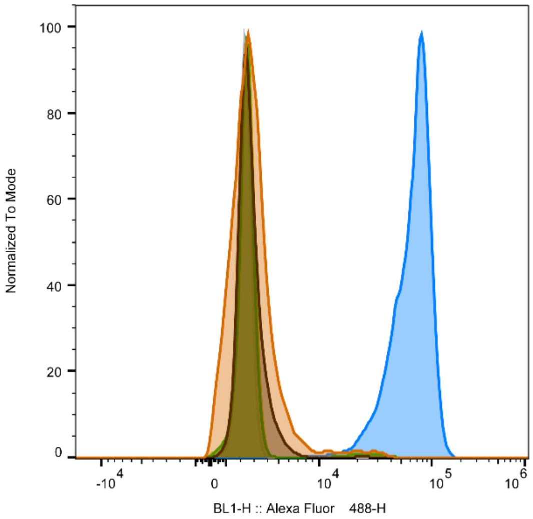


| Bacteria | Media | PyS2-488 |
|---|---|---|
| PAO1 | M9G | |
| PAO1 | M9G | 1 µM |
| PAO1 ΔfpvA1 | M9G | |
| PAO1 ΔfpvA1 | M9G | 1 µM |
| E. coli | M9G | |
| E. coli | M9G | 1 µM |

**Figure S44.** Summary histogram of Figure S40 showing the significant signal difference and development of a distinctly stained PAO1 population in comparison to PAO1 Δ*fpvA* and *E. coli* (ATCC 25922) when grown in iron-poor minimal M9 Glucose (M9G) media.

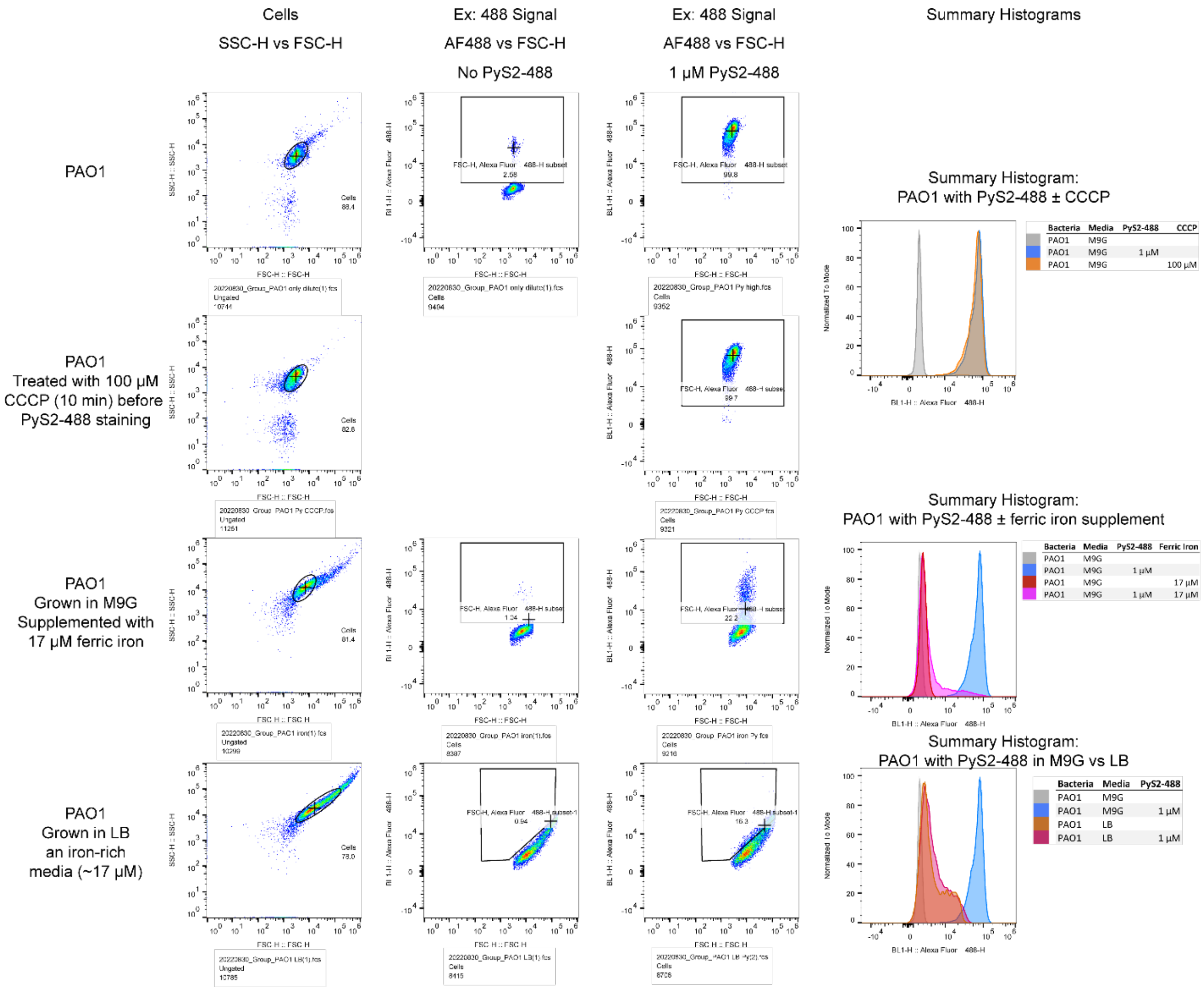


**Figure S42.** Raw representative flow cytometry plots from Main Text Figure 4 indicating FpvA-specific binding of PyS2-488 when grown in iron-poor minimal M9 Glucose (M9G) media in comparison to: i) PAO1 pre-treated with 100 µM CCCP to minimize PyS2-488 internalization but not block fpvA1 binding, ii) PAO1 grown in M9G and supplemented with 17 µM ferric iron (+3) chloride, and iii) PAO1 grown in iron-rich LB media. Summary histograms are shown to the right of each row. 10,000 events were recorded with the final gate number of events listed below each graph. Each graph is a representative of triplicate biologic experiments.

**Table S1.** Geometric means of flow cytometry data shown in Main Text Figure 4 with all biological replicates and their intensities shown upon excitation at 488 nm. Average and standard deviation are calculated for each condition.

| Bacteria | Media | PyS2-488 | CCCP | Ferric Iron (+3) chloride | Geometric Mean, BL1-H: AlexaFluor 488-H | Average ± SD (%SD of Ave) |
|---|---|---|---|---|---|---|
| PAO1 | M9G | | | | 3714 | |
| PAO1 | M9G | | | | 2588 | 2993 ± 626 (20.9%) |
| PAO1 | M9G | | | | 2678 | |
| PAO1 | M9G | 1 µM | | | **61032** | |
| PAO1 | M9G | 1 µM | | | **68733** | **65601 ± 4047 (6.2%)** |
| PAO1 | M9G | 1 µM | | | **67039** | |
| PAO1 ΔfpvA1 | M9G | | | | 3055 | |
| PAO1 ΔfpvA1 | M9G | | | | 2798 | 2857 ± 176 (6.2%) |
| PAO1 ΔfpvA1 | M9G | | | | 2718 | |
| PAO1 ΔfpvA1 | M9G | 1 µM | | | 2674 | |
| PAO1 ΔfpvA1 | M9G | 1 µM | | | 2779 | 2716 ± 56 (2.1%) |
| PAO1 ΔfpvA1 | M9G | 1 µM | | | 2694 | |
| E. coli | M9G | | | | 2990 | |
| E. coli | M9G | | | | 2943 | 2925 ± 76 (2.6%) |
| E. coli | M9G | | | | 2842 | |
| E. coli | M9G | 1 µM | | | 2953 | |
| E. coli | M9G | 1 µM | | | 2901 | 2936 ± 30 (1%) |
| E. coli | M9G | 1 µM | | | 2953 | |
| PAO1 | M9G | 1 µM | 100 µM | | **55399** | |
| PAO1 | M9G | 1 µM | 100 µM | | **53875** | **53870 ± 1532 (2.8%)** |
| PAO1 | M9G | 1 µM | 100 µM | | **52335** | |
| PAO1 | M9G | | | 17 µM | 3149 | |
| PAO1 | M9G | | | 17 µM | 3178 | 3152 ± 25 (0.8%) |
| PAO1 | M9G | | | 17 µM | 3129 | |
| PAO1 | M9G | 1 µM | | 17 µM | 4552 | |
| PAO1 | M9G | 1 µM | | 17 µM | 4322 | 4377 ± 155 (3.5%) |
| PAO1 | M9G | 1 µM | | 17 µM | 4258 | |
| PAO1 | LB | | | | 4527 | |
| PAO1 | LB | | | | 5345 | 5175 ± 582 (11.2%) |
| PAO1 | LB | | | | 5654 | |
| PAO1 | LB | 1 µM | | | 4910 | |
| PAO1 | LB | 1 µM | | | 5524 | 5382 ± 419 (7.8%) |
| PAO1 | LB | 1 µM | | | 5712 | |

#### SI 13.5. Super-resolution microscopy

Stained bacteria were pelleted and resuspended in 1x PBS. Cells were then fixed with 4% formaldehyde for 30 minutes, pelleted, and then resuspended in 0.5% BSA in M9G with 0.05% Tween80 with 5 µM SYTO 40 Blue, and if present, 100 µM CCCP. Each sample was spotted onto a glass-bottomed microwell dish (MatTek Corporation #P35G-1.5-14-C), allowed to settle overnight and covered with a pre-cooled 1% (w/v) agarose pad. Images were collected on an Applied Precision DeltaVision-OMXv4 Super-Resolution Microscope (60x/1.42 NA oil immersion lens, sequential imaging on two sCMOS cameras). SIM reconstruction and warp-based image alignments were performed with Applied Precision softWoRx. Multi-color Image alignments were calculated from an Applied Precision grid test slide, checked with TetraSpeck beads (0.1 μm, Molecular Probes), and verified with an Argo-SIM test slide (Axiom Optics) prior to data collection. Brightness and contrast were identically adjusted with the open-source Fiji distribution of ImageJ. Images were then converted to an RGB format to preserve normalization and then assembled into panels.

**SI 14.0. References**


(1) Hartrampf, N.; Saebi, A.; Poskus, M.; Gates, Z. P.; Callahan, A. J.; Cowfer, A. E.; Hanna, S.; Antilla, S.; Schissel, C. K.; Quartararo, A. J.; Ye, X.; Mijalis, A. J.; Simon, M. D.; Loas, A.; Liu, S.; Jessen, C.; Nielsen, T. E.; Pentelute, B. L. Synthesis of Proteins by Automated Flow Chemistry. *Science* **2020**, *368* (6494), 980–987.

(2) Greenfield, N. J. Using Circular Dichroism Spectra to Estimate Protein Secondary Structure. *Nat. Protoc.* **2006**, *1* (6), 2876–2890.

(3) Greenfield, N. J. Using Circular Dichroism Collected as a Function of Temperature to Determine the Thermodynamics of Protein Unfolding and Binding Interactions. *Nat. Protoc.* **2007**, *1* (6), 2527–2535.

(4) Hmelo, L. R.; Borlee, B. R.; Almblad, H.; Love, M. E.; Randall, T. E.; Tseng, B. S.; Lin, C.; Irie, Y.; Storek, K. M.; Yang, J. J.; Siehnel, R. J.; Howell, P. L.; Singh, P. K.; Tolker-

Nielsen, T.; Parsek, M. R.; Schweizer, H. P.; Harrison, J. J. Precision-Engineering the Pseudomonas Aeruginosa Genome with Two-Step Allelic Exchange. *Nat. Protoc.* **2015**, *10* (11), 1820–1841.

(5) Day, C. A.; Kraft, L. J.; Kang, M.; Kenworthy, A. K. Analysis of Protein and Lipid Dynamics Using Confocal Fluorescence Recovery After Photobleaching (FRAP). *Curr. Protoc. Cytom.* **2012**, *62* (1), 2.19.1-2.19.29.
